# A Shape Sensing Mechanism driven by Arp2/3 and cPLA_2_ licenses Dendritic Cells for Migration to Lymph Nodes in Homeostasis

**DOI:** 10.1101/2022.08.09.503223

**Authors:** Zahraa Alraies, Claudia A. Rivera, Maria-Graciela Delgado, Doriane Sanséau, Mathieu Maurin, Roberto Amadio, Giulia Maria Piperno, Garett Dunsmore, Aline Yatim, Livia Lacerda Mariano, Pablo J. Sáez, Matthieu Gratia, Olivier Lamiable, Aurélie Moreau, Alice Williart, Benoit Albaud, Patricia Legoix, Hideki Nakano, Donald N Cook, Toby Lawrence, Nicolas Manel, Federica Benvenuti, Florent Ginhoux, Hélène D. Moreau, Guilherme P.F. Nader, Matthieu Piel, Ana-Maria Lennon-Duménil

**Author notes:** These authors jointly supervised this work (Guilherme P.F. Nader, Matthieu Piel, Ana-Maria Lennon-Duménil). These authors contributed equally to the experimental part of this work (Maria-Graciela Delgado, Doriane Sanséau). Corresponding authors &.

## Abstract

Motile cells such as immune and cancer cells experience large deformation events that result from the physical constraints they encounter while migrating within tissues or circulating between organs. It has become increasingly clear that these cells can survive and adapt to these changes in cell shape using dedicated shape sensing pathways. However, how shape sensing impacts their function and fate remains largely unknown. Here we identify a shape sensing mechanism that couples cell motility to expression of CCR7, the chemokine receptor that guides immune cells to lymph nodes. We found that this mechanism is controlled by the lipid metabolism enzyme cPLA_2_, requires an intact nuclear envelop and exhibits an exquisitely sensitive activation threshold tuned by ARP2/3 and its inhibitor Arpin. We further show that shape sensing through the ARP2/3-cPLA_2_ axis controls Ikkβ-NFκB-dependent transcriptional reprogramming of dendritic cells, which instructs them to migrate to lymph nodes in an immunoregulatory state compatible with their homeostatic tolerogenic function. These results highlight that the cell shape changes experienced by motile cells evolving within the complex environment of tissues can dictate their behavior and fate.

## Introduction

Cell migration serves many physiological and pathological processes in multicellular organisms, ranging from tissue development to immunity and cancer. To migrate, cells must exert forces on the environment, which in most cases is achieved through the dynamic reorganization of their actomyosin cytoskeleton in response to external signals (1). In addition, to reach a particular destination, cells sense biochemical and/or physical extracellular cues that guide them through the complex environment of tissues and vessels (2). Among them, chemokines have been shown to play a prominent role as they are instrumental to a variety of biological processes, including adaptive immune responses and invasion of healthy tissues by cancer cells (3; 4). Within tissues, chemokines can form gradients that are recognized by specific cell surface receptors. Various types of stimuli can switch on the expression of chemokine receptors, inducing the migration of cells to a specific target or organ. Whether and how cytoskeleton reorganization and chemokine receptor expression occur independently of each other or are integrated within migrating cells in response to environmental cues they encounter remains unclear.

A chemokine receptor that has attracted lots of attention from both immunologists and cancer biologists is CCR7, which recognizes gradients of CCL19 or CCL21 chemokines (5; 6). CCR7 is strictly required for migration to lymph nodes of both cancer and immune cells. In the case of cancer cells, lymph node migration is used as a route for metastatic seeding (7; 8). In the case of immune cells, migration to lymph nodes allows dendritic cells (DCs) to present the antigens they collected in their peripheral tissue of residency to T lymphocytes (9; 10; 11). Antigen presentation in lymph nodes can have two distinct outcomes (12). It can lead to T cell activation when DCs present antigens from a tissue that was inflamed because of infection or tumor growth. Alternatively, it can lead to inactivation of self-reactive T cells, a process referred to as “peripheral immune tolerance” that prevents autoimmune reactions at steady-state, i.e., in the absence of tissue inflammation (13; 14; 11; 15; 16). Unraveling the mechanisms that control CCR7 expression in both immune and cancer cells is therefore critical to understand how the immune system helps multicellular organisms maintain tissue homeostasis and limit tissue infection, tumor growth and metastatic spreading.

The main inducers of CCR7 expression identified so far are microbial components produced by viruses or bacteria and endogenous inflammation mediators such as growth factors and cytokines (10; 17; 18; 19). Interestingly, seminal studies from the Mellman group have shown that mechanical disruption of cell-cell junctions can also induce the expression of CCR7 in DCs (20). However, whether this phenomenon contributes to DC migration to lymph nodes at steady-state remains an open question, as it is unclear whether DCs form such junctions within peripheral tissues. Nonetheless, these results along with others suggest that the physical signals to which DCs are exposed while patrolling peripheral tissues might modify their capacity to express CCR7 and to migrate to lymph nodes in the absence of tissue inflammation (20; 21).

In tissues, a major physical signal experienced by motile cells, including immune and cancer cells, results from the global shape changes and deformation of internal organelles imposed by the physical constrains of their environment. We and others have shown that both immune and tumor cells can adapt and respond to large shape changes when spontaneously migrating through peripheral tissues or dense tumors (22; 23; 24; 25; 26). Such shape changes lead to nuclear deformation events that can activate the lipid metabolism enzyme cytosolic phospholipase 2 (cPLA_2_), a sensor of nuclear envelope stretching (27; 28; 29). Once activated, this enzyme uses phospholipids from the nuclear membrane to produce arachidonic acid (AA) that can further be converted into different lipidic mediators (30; 31; 32; 33). Remarkably, AA production by cPLA_2_ also enhances the contractility of the actomyosin cortex, allowing cells that are physically constrained to release themselves from the traps imposed by dense tissues and keep moving forward (27; 29). Whether the shape changes leading to activation of cPLA_2_ have any additional impact on the ability of DCs or tumor cells to upregulate CCR7, i.e. the chemokine receptor that guides them towards lymph nodes, is unknown.

Here we applied to DCs distinct deformation events of amplitudes that fall within the range of the shape changes they experience *in vivo*. Strikingly, we identified a precise deformation amplitude at which DCs turn on an ARP2/3-cPLA_2_-NFKB-dependent shape sensing mechanism. Activation of this pathway leads to CCR7 upregulation, in addition to enhancing intrinsic DC motility. Accordingly, we found that this signaling axis controls *in vivo* migration of DCs to lymph nodes at steady-state. We further observed that shape sensing imprints DCs with specific immunoregulatory properties, which are compatible with their tolerogenic homeostatic function and distinct from the ones conferred by microbial compounds. We conclude that the interplay between the branched actin cytoskeleton and lipid metabolism enzymes is used by immune cells to build a coordinated response to the physical constrains they encounter, which integrates cell motility with cell chemotaxis and function. These results show that the shape changes experienced by immune cells can dictate their behavior and fate.

## Results

### Co-regulation of cell motility and CCR7 expression in response to cell shape changes

To migrate to lymph nodes and achieve their immune function, DCs must express the CCR7 chemokine receptor, in addition to enhance their intrinsic cell motility. We have previously observed that confinement-induced stretching of the nuclear envelop increases DC motility by promoting actomyosin contractility in a cPLA_2_-dependent manner (27). We hypothesized that such shape change might further trigger CCR7 expression, thereby endowing DCs with the full capacity to reach their next destination. To test this hypothesis, we confined DCs in a controlled manner using a cell confining device (34; 35). Previous observations had revealed that DCs experiencing cell shape changes while migrating in mouse ear skin explants reached minimal diameters of 2 to 4 μm (25). We observed similar deformation events when using intravital microscopy to image the DCs that patrol the dermis of an intact mouse ear *in vivo* (fig. 1a and supplementary movie 1). We therefore chose to monitor CCR7 expression and DC motility upon confinement at 2, 3 and 4 μm heights (fig. 1b).

**Figure 1:**
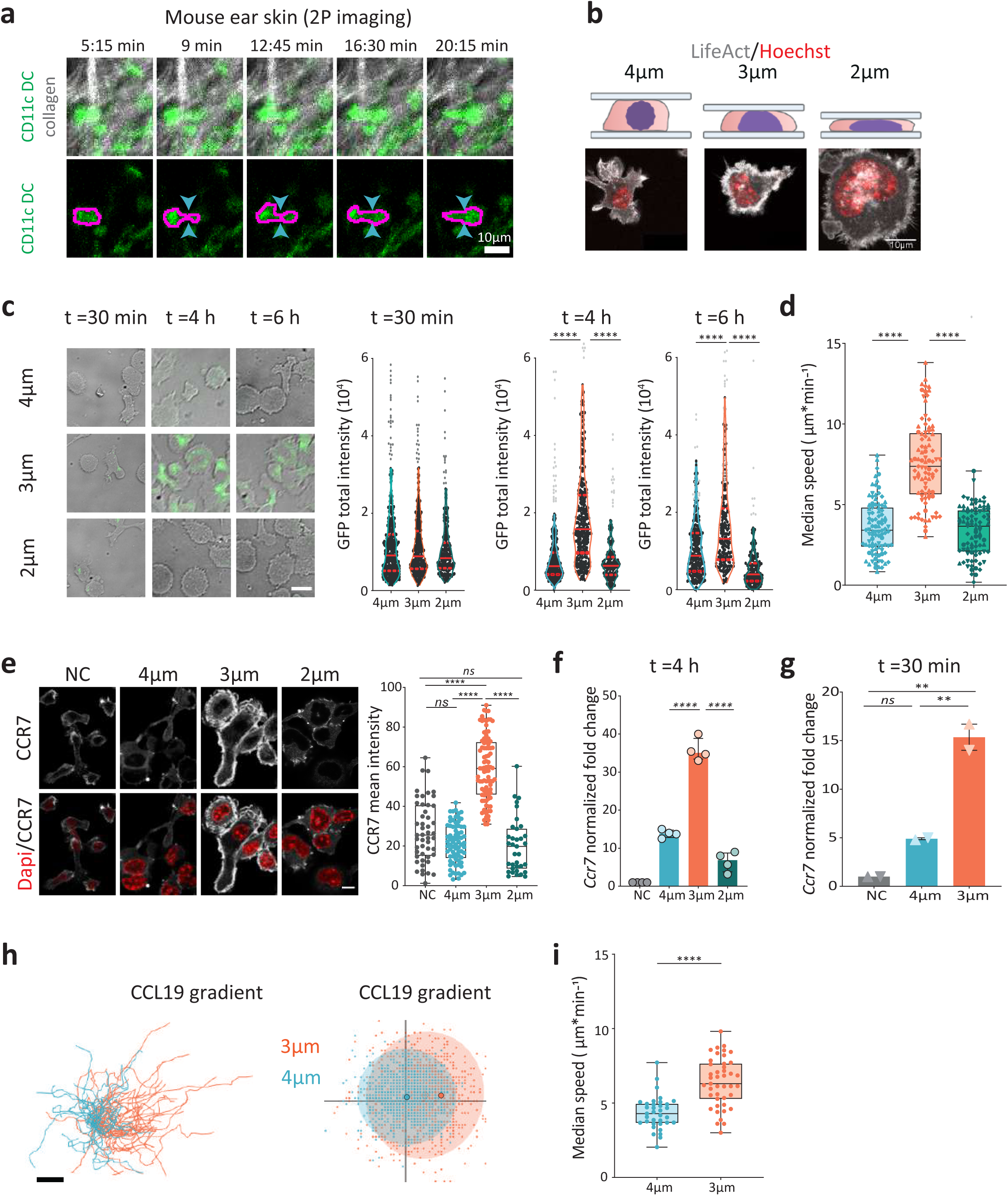
Cell shape sensing can lead to CCR7 upregulation and DC chemotaxis. (**a**) Image sequence from supplementary movie 1. A migrating CD11c-GFP DC imaged by intravital microscopy in the ear dermis. Arrows highlight an example of cell deformation event experienced by the cells during migration. Upper panel: montage of the CD11c-GFP DC with collagen (second harmonic channel) over 5 time points. Lower panel: montage of the GFP channel exclusively. Right part: zoom on one cell passing a constriction of ∼3 µm. Scale bar 10 µm. (**b**) Upper panel: Schematic representation of cells in the different confinement conditions (Createdwith <u>BioRender.com</u>). Lower panel: Representative images of live immature DCs expressing LifeAct-GFP (grey), stained with NucBlue (DNA, red) under different confinement conditions (from left to right: 4, 3, and 2 µm heights). (**c**) Representative images of immature DCs expressing CCR7-GFPunder confinement of 4, 3, and 2 µm heights at different time points upon confinement. Quantification of total GFP intensity in DCs under different confinement heights at various time points. Violin plot representation (medianand quartiles) where each dot is a cell. Outliers were calculated using prism ROUT test(Q=1%) and represented in grey. Left panel: GFP intensity at 30 min of confinement: N=4, n= 348 cells in 4 µm, n=314 cells in 3 µm, n=211 cells in 2 µm. Middle panel: GFP intensity at 4 h of confinement; N=4, n= 201 cells in in 4 µm, n=206 cells in 3 µm, n=176 cells in 2 µm. Right panel: GFP intensity at 6 h of confinement: N=4 n= 215 cells in in 4 µm, n=187 cells in 3 µm), n=180 cells in 2 µm. P value Kruskal-Wallis test ****: p <0.0001. (**d**) Median speed of confined cells at heights of 4, 3, and 2 µm. N=4, n=96 cells in 4 µm, n=89 cells in 3 µm, n=85 cells in 2 µm. P value Kruskal-Wallis test ****: p <0.0001. (**e**) Left panel: Representative immunofluorescence images of immature DCs non-confined or confined for 4 h at 4, 3, and 2 µm. After releasing confinement, CCR7 was visualized using immunostaining (in grey) and the nucleus stained with Dapi (in red). Scale bar of 10 µm.Right panel: Quantification of CCR7 mean intensity, box plot representation where each dot is a cell. N2, n=49 cells in non-confined, n=74 cells in 4 µm, n=89 cells in 3 µm, n=35 cells in 2 µm. P value Kruskal-Wallis ****: p <0.0001, ns: not- significant. (**f**) RT-qPCR data reveals the upregulation of *Ccr7* gene upon 4 h ofconfinement with different heights. Graph: mean with SD of *Ccr7* fold change after normalization to non-confined immature cells. Each dot represents one experiment calculated as described in the methods. N=3, P value Kruskal-Wallis test ****: p <0.0001. (**g**) RT-qPCR data reveals the upregulation of *Ccr7* gene upon 30 min of confinement with different heights. Cells were left in the incubator for 4 h after confinement was removed, and then collected for qPCR. Mean with SDof *Ccr7* fold change after normalization to non-confined control cells. Each dot represents one experiment calculated as described in the methods. N=2, P value Kruskal-Wallis test **: p=0.0021 (A-C), **: p=0.0052 (B-C). (**h-i)** Chemotactic response of DCs confined at 4 (blue) and 3 (orange) µm heights towards a CCL19 gradient, representative of 2 independent experiments, n= 33 cells in 4 µm n= 43 cells in 3 µm. (**h**) Left panel: Cell trajectories (gradient from right to left). Scale bar 100 µm. Middle panel: Representation of each step performed by the cells: the center of the 3 µm cells’ tracks is shifted towards the right,indicating an increase in the directionality of these cells towards the gradient. (**i**) Quantification of the mean speed of DCs in the presence of the CCL19 gradient, box plot representation where each dot is a cell, P value Mann-Whitney test ****: p <0,0001.

As a cell model, we focused on unstimulated bone-marrow-derived DCs, also referred to as “immature DCs”, which express low levels of CCR7 in culture (36; 37), similarly to DCs patrolling peripheral tissues (38). Of note, tissue-resident DCs cannot be used for our purpose as they spontaneously upregulate CCR7 expression upon tissue disruption and cell purification. Bone marrows were obtained from a mouse knocked-in for a CCR7/GFP reporter gene to evaluate by live imaging the expression dynamics of the chemokine receptor (39). Strikingly, fluorescence quantification showed that while GFP expression increased in DCs confined at 3 µm height for 4-6h (supplementary fig. S1a), it was not significantly modified in cells confined at 2 nor at 4 µm-height (fig. 1c, d and supplementary movie 2). GFP upregulation did not result from cell death, which exhibited low rates at the three confinement heights (< 8% of death in all confinement conditions, supplementary fig. S1b). A similar sensitivity window was observed when monitoring the migration speed of confined DCs that increased at 3 but not at 2 nor at 4 µm confinement height (fig. 1d). These experiments indicate that CCR7 expression and cell motility are co-regulated in response to precise cell shape changes, suggesting that these events might indeed be mechanistically coupled.

These results were confirmed analyzing the endogenous CCR7 protein by immunofluorescence and CCR7 mRNA levels by real time quantitative RT-PCR analyses: only immature DCs confined at 3 μm- height exhibited a significant increase in CCR7 expression with no significant change upon confinement at 4 μm-height (fig. 1e, f). We therefore used 4 μm-height as a control height of confinement in all following experiments. The specificity of the anti-CCR7 antibody was verified using CCR7 knock out DCs (supplementary fig. S1c). These results indicate that GFP upregulation upon confinement at 3 µm height did not result from artefactual expression of the knocked in *GFP* gene. Of note, *Ccr7* expression was also induced when cells were confined for 30 min, harvest and analyzed 4h later by quantitative RT-PCR (fig. 1g), suggesting that imposing a 30 min-deformation at 3μm-height on unstimulated DCs was sufficient to upregulate *Ccr7* expression. We further observed that CCR7 expressed at the surface of DCs confined at 3 µm-height was fully functional: addition of its ligand CCL19 at one side of the confinement device led to chemotaxis of DCs confined at 3 but not of cells confined at 4 µm-height (fig. 1h, 1i). Altogether, these results show that confinement of immature DCs at 3, but not 2 nor 4 μm- height leads to up-regulation of CCR7 expression, endowing these cells with the ability to perform chemotaxis in addition to enhancing their intrinsic motility.

### CCR7 upregulation upon shape sensing relies on cPLA_2_ accumulation in the nucleus of DCs

Our results showing that both CCR7 expression and DC motility were induced at the same confinement height (3 μm) suggested that these processes might be controlled by common mechanisms. They prompted us to investigate the involvement of cPLA_2_, as we had previously shown that this lipid metabolism enzyme promotes cell motility in response to nuclear deformation (27; 29). We found that cPLA_2_ knock down (supplementary fig. S2a) abrogated the upregulation of CCR7/GFP expression observed in DCs confined at 3 µm-height and decreased their increase in motility (fig. 2a, supplementary fig. S2b), 4 µm confined cells were used as a control. To strengthen these results, we generated conditional knock out mice for the cPLA_2_ gene (*Pla2g4a*^flox/flox^ crossed to CD11c-CRE transgenic animals to obtain cPLA_2_ DCs). We found that these cells did not upregulate CCR7 expression nor increase their motility upon confinement at 3 µm-height (fig.2b, c). In sharp contrast, cPLA_2_ knock down or knock out did not prevent the upregulation of CCR7 expression in response to treatment with the microbial compound lipopolysaccharides LPS (supplementary fig. S2c, d). These results indicate that cPLA_2_ is specifically required for CCR7 expression induced by cell shape changes, rather than being generally involved in the transcriptional regulation of the *Ccr7* gene. We conclude that upregulation of both cell motility and CCR7 expression in DCs confined at 3 µm-height rely on the activity of the cPLA_2_ enzyme.

**Figure 2:**
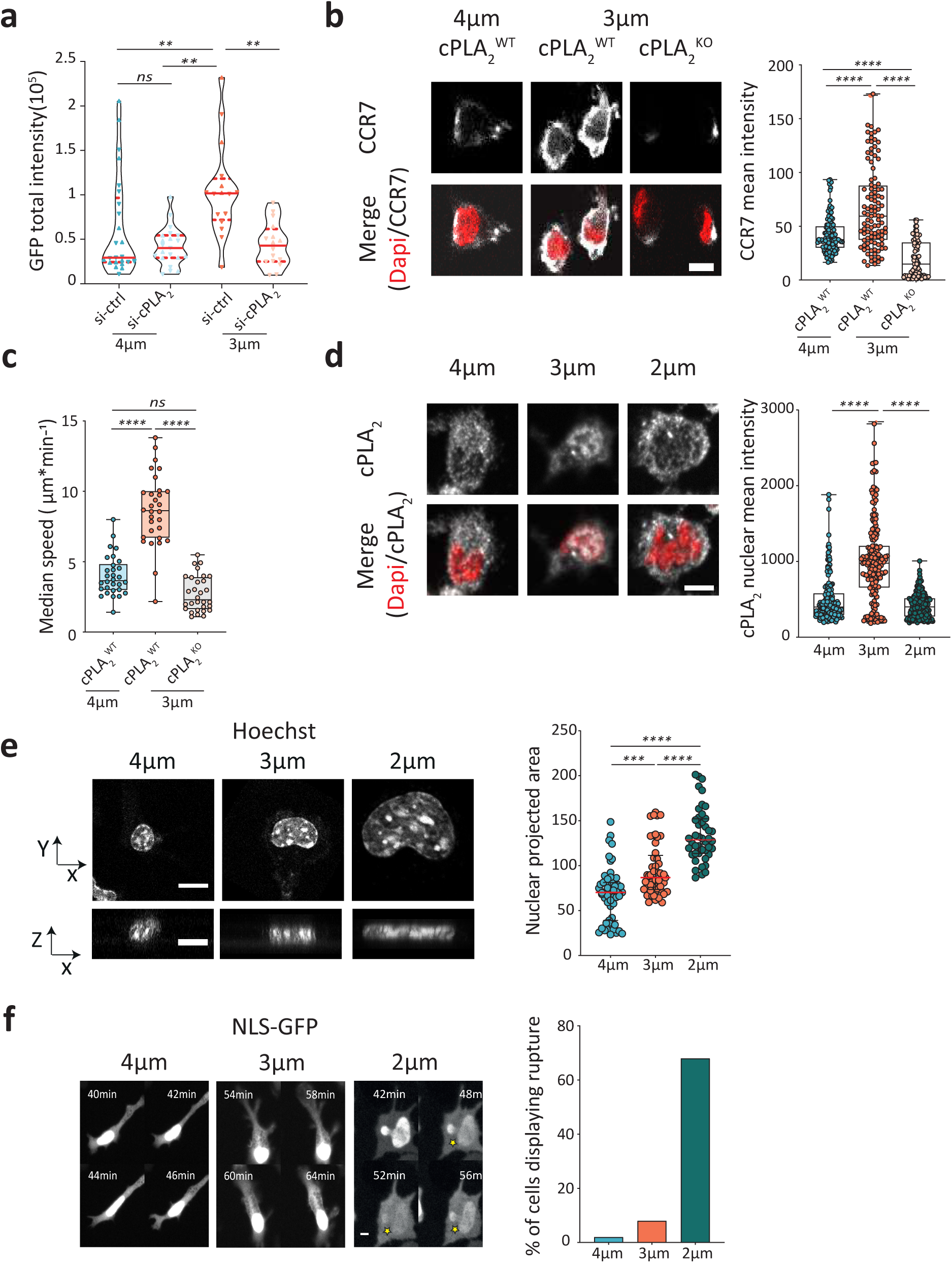
CCR7 upregulation in response to shape sensing depends on cPLA_2_ and requires an intact nuclear envelope. **(a)** Quantification of total GFP intensity in DCs from CCR7-GFP mice after knocking down or not cPLA_2_ using si-RNA and confining cells for 4 h. Violin plot (median and quartiles), each dot is a cell, N=2, n= 25 cells in 4 µm si-ctrl (80% of cells), n=21 cells in 4 µm si-cPLA_2_ (75% of cells), n= 16 cells in 3 µm si- ctrl (75% of cells), n=16 cells in 3 µm si-cPLA_2_ (76%of cells), P value Kruskal-Wallis test **: p= 0,0059 (A-C), P= 0,0039. (**b**) Left panel: Representative immunofluorescence images of DCs from cPLA_2_^WT^ (*flox/flox Cre-*) and KO (*flox/flox Cre+*) mice confined at 4 and 3µm. CCR7 in grey and the nucleus in red. Scale bar of 10 µm.Right panel: Quantification of CCR7 mean intensity, box plot representation where each dot is a cell. N=2, n=114 cells in 4 µm WT, n=105 cells in 3 µm WT, n= 86 cells in 3 µm KO. P value Kruskal-Wallis ****: p <0.0001. (**c**) Median speed of confined cells from cPLA_2_ WT and KO at heights of 4 and 3 µm. N=2, n30 6 cells in all positions. P value Kruskal-Wallis test ****: p <0.0001. (**d**) Left part: representative immunofluorescence images of immature DCs confined for 4 h. cPLA_2_ is shown in grey and the nucleus in red. Scale bar of 10 µm. Right part: Quantification of cPLA_2_ mean intensity, box plot representation where each dot is a cell. N=3, n=165 cells in 4 µm, n=178 cells in 3 µm, n=230 cells in 2 µm, P value Kruskal-Wallis test ****: p <0.0001. (**e**) Left panel: Nucleus of cells under confinement of 4,3, and 2 µm represented in different axes. Scale bar 3 µm. Right panel: Quantification of the nucleus projected area, box plot representation where each plot is a cell. N=2, n=55 cells in 4 µm, n=49 cells in 3 µm, n=44 cells in 2 µm. P value Kruskal-Wallis test ****: p <0.0001, ***: p <0.0006 (**f**) Left part: representative images of DCs transduced with NLS-GFP (GFP signal in grey) to visualize nuclear envelope rupture in response to confinement at 4, 3, and 2 µm-heights. Yellow stars were drawn to indicate nuclear envelop rupture and repair events. Right part: quantification of the percentage of DCs displaying rupture events during the first hour of confinement at different heights, N=2.

In good agreement with these data, we observed that cPLA_2_ accumulated in the nucleus of DCs confined at 3 but not 2 nor 4 µm-heights (fig. 2d). Indeed, this enzyme was shown to translocate to the nucleus (40) and accumulate at the inner nuclear membrane upon activation (41; 42). Of note, analysis of nuclear shape showed that DCs confined at both 3 and 2 μm-heights gradually increased their nucleus projected area (fig. 2e), consistent with their nuclei being more deformed than the nuclei of non-confined or 4 µm-height confined cells. Thus, cPLA_2_ does not accumulate into the nucleus of DCs confined at 2 µm-height despite their nucleus being extensively stretched. These results prompted us hypothesizing that confinement at 2 µm-height might compromise nuclear accumulation of cPLA_2_ due to loss of nuclear envelope integrity. To test this hypothesis, we transduced DCs with a lentiviral construct expressing nuclear localization signal (NLS)-GFP. We observed that most NLS-GFP-expressing DCs confined at 2 µm-height underwent events of nuclear envelope rupture followed by repair, as evidenced by the transient leakage of NLS-GFP signal into their cytoplasm (fig. 2f and supplementary movie 3). Nuclear envelope rupture was less frequently observed in DCs confined at 3 or 4 µm-height (fig. 2f). Of note, DCs confined at 2 µm-height did not display any additional sign of damage and were able to upregulate *Ccr7* expression upon treatment with the microbial compound lipopolysaccharide (LPS) (supplementary fig. S2e). Altogether these results show that coordinated upregulation of DC motility and CCR7 expression relies on nuclear accumulation of cPLA_2_, which requires an intact nuclear envelope. They point to DCs being equipped with an extremely accurate machinery to detect precise levels of cell shape changes.

### The cPLA_2_ activation threshold to cell shape changes is defined by ARP2/3 and its Arpin inhibitor

We next asked whether nuclear translocation and activation of cPLA_2_ resulted from passive stretching of the DC nucleus upon confinement or rather required an active cellular response. A good candidate for driving such response was ARP2/3, as this complex had been shown to nucleate distinct types of actin structures in the perinuclear area of DCs undergoing nucleus deformation (43; 23). Live-imaging analysis of LifeAct-GFP distribution in DCs confined at 3 µm-height indeed revealed a cloud of perinuclear F-actin that was not observed in cells confined at 4 µm. Of note, this actin cloud was not always observed in DCs unconfined, fixed, and stained with phalloidin, suggesting that it could be lost upon deconfinement and/or fixation. Treatment of DCs with CK666, which inhibits ARP2/3 activity led to disappearance of this actin structure (fig. 3a, supplementary movie 4, and supplementary fig. S3a for quantification), suggesting that ARP2/3 might be involved in the DC response to shape changes. Accordingly, we found that CK666 impaired the upregulation of *Ccr7* expression and the nuclear accumulation of cPLA_2_ upon confinement at 3 µm-height (fig. 3b, c). As observed for cPLA_2_, ARP2/3 inhibition had no effect on LPS-induced CCR7 upregulation (supplementary fig. S3b). Consistent with the involvement of ARP2/3 in the cell response to shape changes, we further found that DCs knocked out for WASp (Wiscott Aldrich Syndrome Protein), which was recently shown to activate ARP2/3 in the DC perinuclear area (44), behaved as cells treated with CK666: they did not upregulate CCR7 expression nor showed cPLA_2_ nuclear translocation upon confinement at 3 µm-height (fig. 3d,e). Altogether these results suggest that the response of DCs to shape changes requires WASp and ARP2/3 activity to allow nuclear accumulation of cPLA_2_ and subsequent upregulation of CCR7 expression.

**Figure 3:**
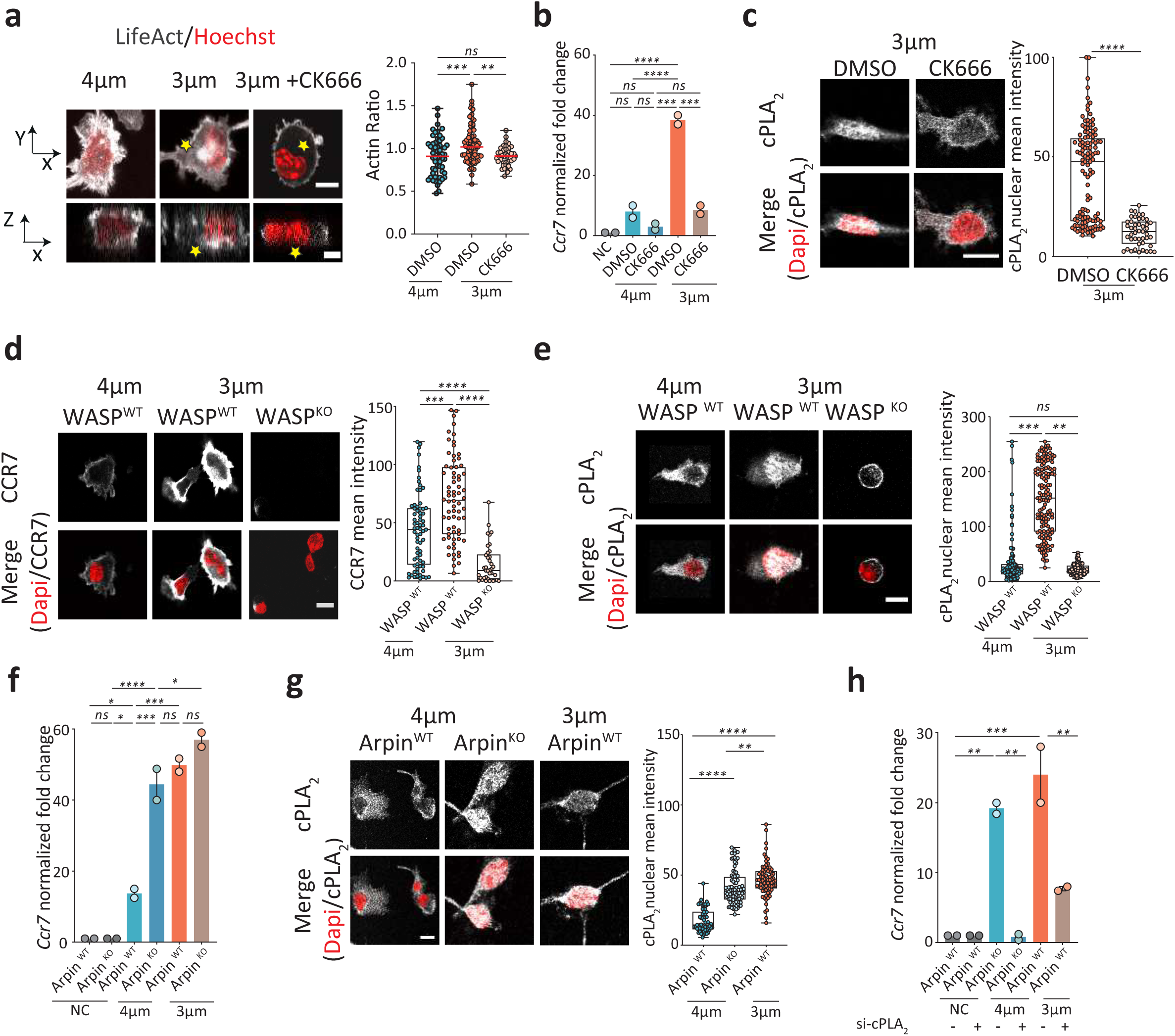
Arp2/3 activity tunes the sensitivity of the cPLA_2-_dependent shape sensing DC response. (**a**) Left part: representative images of DCs expressing LifeAct-GFP (grey) stained with NucBlue (DNA, red) under confinement of 4 and 3 µm heights and treated with CK666 (25µM) or DMSO. Yellow stars indicate the actin cloud in the perinuclear area. Right part: Quantification of LifeAct perinuclear to cytosolic ratio calculatedas explained in supplementary figure 3. N=2, n=55 cells in 4 µm DMSO, n=70 cells in 3 µm DMSO, n=42 cells in 3 µm CK666. P value ordinary one-way ANOVA test ***: p=0.0007, **: p=0.0016. (**b**) *Ccr7* gene expression upon 4 h of confinement of cells treated with CK666 (25µM) or DMSO, as measured by RT-qPCR. Mean with SD of *Ccr7* fold change after normalization to non- confined control cells. Each dot represents one experiment. N=2, P value ordinary one-way ANOVA test ****: p <0.0001, ***: p =0.0001. (**c**) Left part: representative immunofluorescence images of immature DCs treated either with CK666(25 µM) or DMSO and confined for 4 h at 3 µm-height. cPLA_2_ is shown in grey and the nucleus in red. Scale bar of 10 µm. Right part: Quantification of cPLA_2_ mean intensity, box plot representation where each dot is a cell. N=2, n=121 cells in DMSO-3 µm, n=64 cells in CK666-3µm, P value Mann-Whitney test p <0.0001. (**d**) Left panel: representative immunofluorescence images of immature DCs from WASp WT and KO mice confined for 4 h. CCR7 is shown in grey, the nucleus in red. Scale bar of 10 µm. Right panel: quantification of CCR7 mean intensity, box plot representation where each dot is a cell. N=2, n=70 cells in 4 µm WT n=70 cells in 3 µm WT, n=38 cells in 3 µm KO, P value Kruskal-Wallis test ****: p <0.0001 ***: p 0,0002. (**e**) Left panel: representative immunofluorescence images of immature DCs from WASp WT and KO mice confined for 4h. cPLA_2_ is shown in grey and the nucleus in red. Scale bar of 10 µm. Right panel: Quantification of cPLA_2_ mean intensity, box plot representation where each dot is a cell. N=2, n=150 cells in 4 µm-WT n=145 cells in 3 µm-WT, n=149 cells in 3 µm-KO, P value Kruskal-Wallis test ****: p <0.0001. (**f**) RT-qPCR data of *Ccr7* gene expression upon 4 hour-confinement of cells from Arpin WT and KO mice. The graph represents the mean with SD of *Ccr7* fold changes observed after normalization to non-confined control cells. Each dot represents one experiment. N=2, P values ordinary one-way ANOVA test ***: p = 0.0002, **: p= 0.0015. (**g**) Left panel: representative immunofluorescence images of immature DCs from Arpin WT and KO mice confined for 4h. cPLA_2_ is shown in grey and the nucleus in red. Scale bar of 10 µm. Right panel: Quantification of cPLA_2_ mean intensity, box plot representation where each dot is a cell. N=2, n=59 cells in WT-4 µm, n=87 cells in KO-4 µm, n=75 cells in WT-3 µm, P value ordinary one-way ANOVA test****: p <0.0001, **: P= 0.0043. (**h**) RT-qPCR data of *Ccr7* gene expression upon 4h of confinementof DCs from Arpin^WT^ or Apin^KO^ that have been knocked down for cPLA_2_ using siRNA (as in Figure 2). The graph represents the mean with SD of *Ccr7* fold changes observed after normalization to non-confined is showncontrol cells. Each dot represents one experiment. N=2, P values ordinary one-way ANOVA test **: p = 0.0046 (B-C), **: p= 0.0080.

To strengthen these findings, we assessed the response of DCs knocked out for the ARP2/3 inhibitor Arpin (*Arpin ^f^*^lox/flox^ x CD11c-CRE), which exhibit enhanced ARP2/3 activity (45). Remarkably, we found that Arpin^KO^ immature DCs displayed an increased sensitivity to cell shape changes as they upregulated *Ccr7* when confined at 4 instead of 3 µm-height (fig. 3f). In addition, Arpin^KO^ cells confined at 4 µm exhibited levels of cPLA_2_ nuclear translocation like those found in Arpin^WT^ DCs confined at 3 µm-height (fig. 3g). Induction of *Ccr7* expression in Arpin^KO^ DCs confined at 4 µm-height was also cPLA_2_-dependent (fig. 3h). We conclude that the activity of ARP2/3 determines the sensitivity of DCs to confinement, defining a threshold for cPLA_2_ nuclear accumulation and induction of CCR7 expression in response to cell shape changes. These results are consistent with DCs being equipped with a sensory mechanism that finely tunes their response to shape changes.

### ARP2/3-cPLA_2_-dependent shape sensing drives DC migration to lymph nodes at steady-state

So far, we have shown that WASp-ARP2/3-dependent actin remodeling in response to shape changes tunes the activation threshold of cPLA_2_ and thereby defines the capacity of DCs to upregulate the two elements required for migration to lymph nodes: increased intrinsic cell motility and CCR7 expression. These data strongly suggest that (1) WASp-ARP2/3-dependent cPLA_2_ activation in response to cell shape changes might license DCs for migration to lymph nodes in the absence of inflammation (due to infection or tumor growth) and (2) by restraining the activation of this shape sensing pathway, Arpin might act as a negative regulator of this process *in vivo*. Such regulatory mechanism could limit the number of DCs that become CCR7-positive and thus migrate to lymph nodes upon the shape changes they experience while patrolling peripheral tissues, thereby increasing their time of tissue-residency. To assess the *in vivo* physiological relevance of shape sensing by ARP2/3 and cPLA_2_, we evaluated by flow cytometry whether cPLA_2_, WASp and Arpin deficiencies altered the number of migratory DCs present in skin-draining lymph nodes at steady-state.

Skin DCs can be divided in two main subtypes based on surface markers: conventional DCs type 1 and type 2 (46; 47; 48). Although both populations can migrate to lymph nodes, the migration rates of cDC2s have been found to be more elevated than those of cDC1s at steady-state (46). Strikingly, we observed that the numbers of migratory cDC2s found in inguinal lymph nodes were significantly decreased in WASp^KO^ and cPLA_2_^KO^ mice (fig. 4a, b, c). No such difference was detected for migratory cDC1s, which, as expected from other’s findings, were less represented in these secondary lymphoid organs in homeostatic conditions. Conversely, we found that Arpin^KO^ mice displayed enhanced numbers of migratory cDC2s in lymph nodes as compared to their wild-type counterpart (fig.4 d). Of note, analysis of DC numbers in the skin of these animals showed no significant difference (fig.4 e, f, g), excluding that the differences observed in lymph node cDC2 numbers could result from altered cDC2 development and/or survival in the skin of WASp^KO^, cPLA_2_^KO^ or Arpin^KO^ mice. Hence, ARP2/3 activity controlled by WASp and Arpin finely tunes the number of DCs that migrate to lymph nodes at steady-state, possibly by controlling the cPLA_2_ activation threshold and downstream CCR7 expression in response to nucleus deformation. These data suggest that steady-state migration of DCs to lymph nodes might be dictated at least in part by the events of shape changes they experience while patrolling the complex environment of the skin.

**Figure 4:**
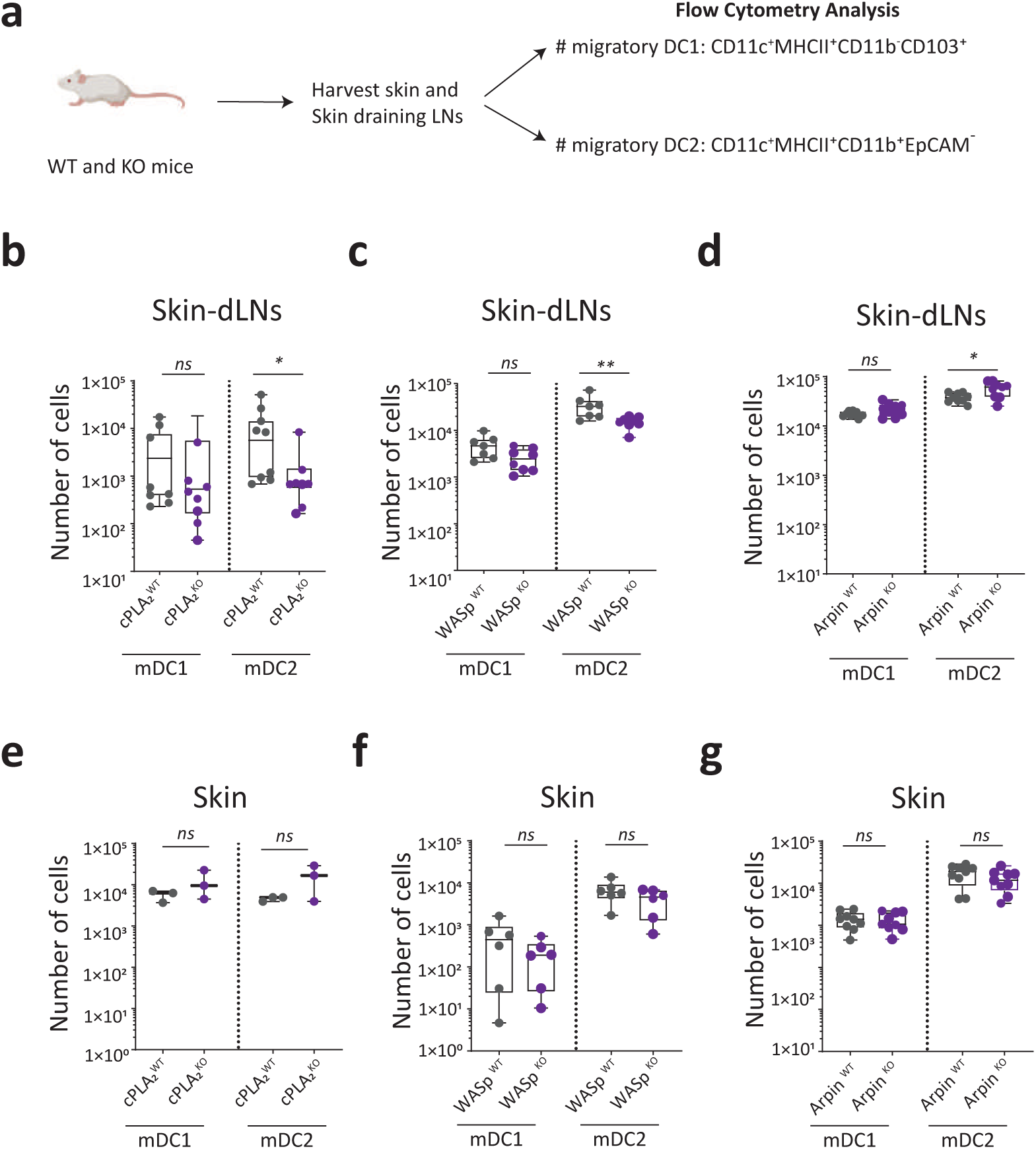
Steady-state migration of skin mDC2 is cell shape sensitive. **(a)** Gating strategy to quantify DCs in skin-draining lymph nodes of mice at steady-state: after gating on live cells, immune cells were identified as CD45^high^; CD11c and MHCII were then used to differentiate lymph node-resident DCs (MHCII^low^, CD11c^high^) from migratory DCs (MHCII^high^, CD11c^high^). Among migratory DCs, mDC1s were identified as CD11b^low^, CD103^high^ and mDC2s as CD11b^high^, EPCAM^low^. (**b**) Plots inLogscaleof thenumber of migratoryDCs in the inguinal skin-draining lymph nodes of cPLA_2_ (*flox/flox Cre-*) and KO (*flox/flox Cre+*) mice, N=3 where each dot is a mouse. P value Mann- Whitney test *: p= 0,0221, ns: non-significant. (**c**) Plots in Log scale of number of migratoryDCs in the inguinal skin draining lymph nodes of WASp^WT^ and WASp^KO^ mice (total KO), N=3 where each dot is a mouse. P value Mann-Whitney test **: p= 0,0059, ns: non-significant. (**d**) Plots in Logscaleof number ofmigratoryDCs in the inguinal skin draining lymph nodes of Arpin^WT^ (*flox/flox Cre-/-*) and KO (*flox/flox Cre+/-*) mice, N=3 where each dot is a mouse. P value Mann-Whitney test *: p= 0,04, ns: non- significant. (**e**) Plots inLogscaleshowing the number of DCs in the skin of cPLA_2_ and cPLA_2_ mice, N=1 where each dot is a mouse. P value Mann-Whitney test *: p= 0,0221, ns: non-significant. (**f**) Plots in Logscale showing the number of migratoryDCs in the skin of WASp ^WT^ and WASp ^KO^ mice, N=2 where each dot is a mouse. P value Mann-Whitney test, ns: non-significant. (**g**) Plots in Logscale showing the number of migratoryDCs in the skin of Arpin ^WT^ and KO mice, N=3 where each dot is a mouse. P value Mann-Whitney test, ns: non-significant.

### The ARP2/3-cPLA_2_ shape sensing axis controls DC steady-state migration by activating Ikkβ and NFκB

The cDC2s that migrate from the skin to lymph nodes at steady-state were shown to display a specific transcriptional profile enriched for NFκB- and type I interferon (IFN)-related genes (46). NFκB activation in response to the kinase Ikkβ (*Ikbkb*) was further described as required for migration of DCs from the skin to lymph nodes at steady-state and upon inflammation (49). Of note, this pathway is the only one described so far as implicated in homeostatic DC migration to these lymphoid organs. We therefore investigated whether the ARP2/3-cPLA_2_ shape sensing axis here identified as leading to DC homeostatic migration requires the Ikkβ-dependent NFκB activation and whether it also triggers a more profound transcriptional reprogramming of DCs. To this mean, we compared by bulk RNAseq the transcriptome of cPLA_2_^WT^ and cPLA_2_^KO^ DCs confined at 3 µm-height; non-confined cells were used as negative controls.

Principal component and clustering analyses revealed that while non-confined non-stimulated cPLA_2_^WT^ and cPLA_2_^KO^ (NC NS cPLA_2_^WT/KO^) samples clustered together, this did not apply to confined cPLA_2_^WT^ and cPLA_2_^KO^ cells, showing that they display important differences in their gene expression profiles (fig. 5a). These results indicate that cPLA_2_ impacts on the transcriptome of confined DCs but has no major effect on non-confined cells at steady-state. More specifically, we observed that ∼5000 and ∼4600 genes were respectively up- and down-regulated in wild-type DCs confined at 3 µm-height compared to non-confined condition (fig. 5b). Comparison of cPLA_2_^WT^ and cPLA_2_^KO^ DCs showed that more than half of the genes upregulated by confinement relied on cPLA_2_ (∼3600 genes, fig. 5b). Consistent with our findings, *Ccr7* upregulation was found to be fully lost in cPLA_2_^KO^ DCs, which even showed a decrease in *Ccr7* mRNA levels upon confinement (fig. 5c). Strikingly, among the 103 genes following the same expression pattern than *Ccr7* were two genes associated with ARP2/3-dependent actin nucleation (*Actr2*, *Actr3*), the Ikkβ gene (*Ikbkb*) itself and several IFN genes (ISGs) (fig. 5d). Direct comparison of the transcriptional profiles of our confined bone-marrow-derived DCs with the one described for skin migratory cDC2s revealed that they exhibited similar signatures (for the genes following *Ccr7* expression pattern, fig. 5e). These findings therefore highlight that the ARP2/3-cPLA_2_ shape sensing axis imprints DCs with a similar transcriptional program than the one displayed by skin cDC2s migrating to lymph nodes at steady-state.

**Figure 5:**
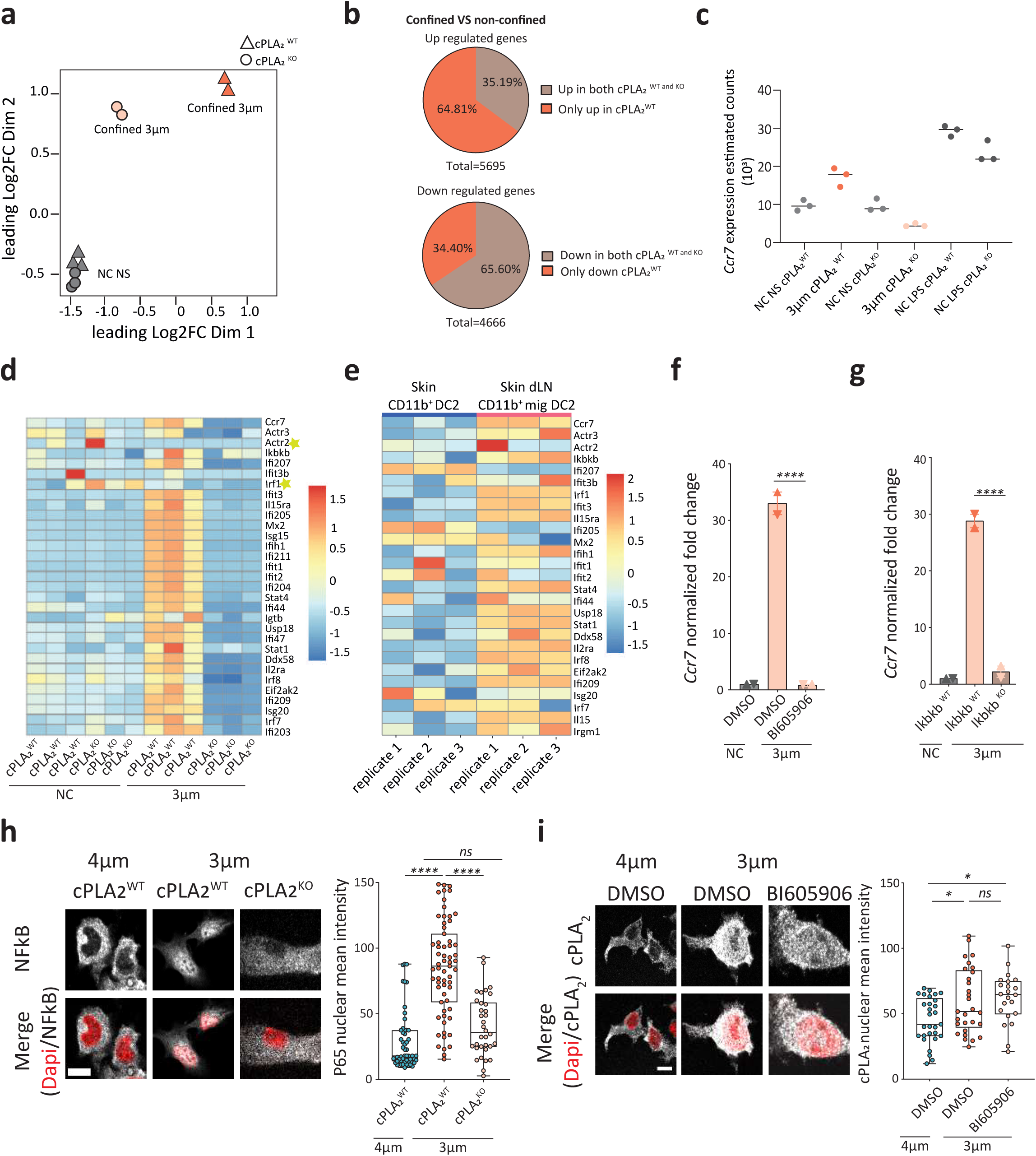
cPLA_2_ reprograms DC transcription in an *Ikk*β-NFκB-dependent manner. (**a-d**) Bulk RNA-seq analysis of BMDCs cPLA_2_^WT^ (*flox/flox Cre-*) and KO (*flox/flox Cre+*) mice. (**a**) ^Data^ from bulk RNA-seq analysis on BMDCs from cPLA2 WT and cPLA2 KO mice. (**a**) Multidimensional scaling (MDS) of the bulk RNA-Seq samples. The MDS figure represents a two-dimensional scatterplot of the first two principal components of the RNA- Seq data. Sample groups at different confinement conditions are represented by different forms. Each dot represents a biological replicate of an RNA- Seq sample (**b**) Pie charts showing proportions of differentially expressed genes in confined conditions compared to non-confined (FDR < 0,05, and Log2FC < -1.0 or > 1.0). Upper part: number of up regulated genes in response to confinement in both cPLA2 WT and KO DCs (5695 genes) in which 3691 of them are up regulated in cPLA2WT compared to KO confined DCs. Lower part: number of down regulated genes in response to confinement in both cPLA2 WT and KO DCs (4666 genes) in which 1605 of them are down regulated in cPLA2WT compared to KO confined DCs. (**c**) *Ccr7* estimated gene counts in the different samples/conditions. (**d**) Heat map of genes harboring a similar expression pattern than *Ccr7*. (**e**) Heatmap showing normalized expression of some ISG genes from **d** in the CD11b+ DC2 subset from dermis and their migratory counterparts in the cutaneous draining lymph node of healthy mice (GSE49358 microarray study from Tamoutounour et al., 2013). (**f**) RT-qPCR data of *Ccr7* gene expression upon 4h of confinement of cells treated with Ikkβ inhibitor BI605906 (30 µM) or with DMSO. The graph represents the mean with SD of *Ccr7* fold change after normalization to non-confined control cells. Each dot represents one experiment. N=2, P values ordinary one- way ANOVA test ****: p<0.0001, *: p= 0.0134. (**g**) RT-qPCR data of *Ccr7* gene expression upon 4hof confinement in cells from *Ikbkb* ^WT^ or *Ikbkb* ^KO^ mice that were confined at the height of 3 µm. The graph represents the mean with SD of *Ccr7* fold change after normalization to non-confined control cells. Each dot represents one experiment. N=2, P values ordinary one-way ANOVA test ****: p<0.0001, ***: p= 0.0001, ns: non-significant. (**h**) Left panel: representative immunofluorescence images of DCs from cPLA_2_ WT and KOs confined for 4h. NFκ_2_B (P65) is shown in grey and th_2_ e nucleus in red. Scale bar of 10 µm. Right panel: Quantification of NFκB (P65) mean intensity, box plot representation where each dot is a cell. N=2, n=55 cells in cPLA_2_ ^WT^-4 µm, n=63 cells in cPLA_2_ ^WT^-3 µm, n=34 cells in cPLA_2_ ^KO^-4 µm, P value Kruskal-Wallis test ****: p <0.0001, ns= non-significant. (**i**) Left part: representative immunofluorescence images of immature DCs treated with Ikkβ inhibitor BI605906 (30µM) or with DMSO and confined for 4h at the height of 3 µm. cPLA_2_ is shown in grey and the nucleus stained in red. Scale bar of 5 µm. Right panel: Quantification of cPLA_2_ mean intensity, box plot representation where each dot is a cell. N=2, n=29 cells in DMSO, n=23 cells in BI605906, P values ordinary one-way ANOVA test *: 0.0175, ns: non-significant.

These results prompted us to investigate whether cPLA_2_ and Ikkβ-NFκB were part of the same signaling pathway. To address this question, we first assessed whether Ikkβ was needed for CCR7 upregulation upon shape sensing in DCs. Remarkably, we found that confinement at 3 µm-height did not lead to *Ccr7* upregulation in DCs treated with the Ikkβ inhibitor BI605906 or knocked out for the *Ikbkb* gene (fig. 5f, g). Consistent with this result, we observed that NFκB nuclear translocation was compromised in confined DCs lacking cPLA_2_ (fig.5h). In contrast, Ikkβ inhibition had no effect on nuclear accumulation of cPLA_2_ in confined DCs (fig.5i). These data strongly suggest that cPLA_2_ acts upstream of Ikkβ and NFκB nuclear translocation to trigger upregulation of CCR7 expression upon shape sensing in DCs. Altogether our results support a model where shape sensing through the ARP2/3-cPLA_2_ axis activates the Ikkβ- NFκB pathway and thereby licenses DCs to migrate to lymph nodes at steady-state.

### The ARP2/3-cPLA_2_ shape sensing axis endows DCs with specific immunoregulatory properties

DC migration to lymph nodes at steady-state helps maintaining peripheral tolerance by transmitting tolerogenic signals to the T lymphocytes that recognize antigens from self (5; 13; 11; 50; 51). This is in sharp contrast to microbe-induced DC migration to lymph nodes that leads to activation of T cells capable of fighting these infectious agents. Our results showed that ARP2/3 and cPLA_2_ were required for CCR7 upregulation in response to confinement, but not in response to LPS (fig. 5c). This suggests that the shape sensing pathway here described might be specifically involved in steady-state rather than microbe-induced migration of DCs. This scenario would be particularly appealing as no specific mechanism has been identified so far for the triggering of homeostatic DC migration, the *Ikbkb* -NFκB pathway being required for both DC migration to lymph nodes at steady-state and upon inflammation (49). To test this hypothesis, we further analyzed the transcriptomes of DCs expressing or not cPLA_2_, either stimulated by confinement at 3 µm-height or by a microbial component (LPS). Strikingly, principal component and clustering analyses of non-confined DCs treated or not with LPS revealed that cPLA_2_ had no significant impact on the global gene expression pattern induced by microbial stimulation (fig.6a, supplementary fig. S4a), consistent with our hypothesis. Of note, we observed that the expression of some genes from the cPLA_2_ pathway were increased in both DCs confined at 3 µm-height and DCs activated with LPS as compared to non-confined non stimulated cells. Yet, the increase in cPLA_2_ and prostaglandin receptor gene expression were more pronounced in confined DCs as compared to LPS-treated cells (fig. 6b). Altogether these data strongly suggest that shape sensing reprograms DC transcription in a way distinct from microbial activation and with a specific requirement for the cPLA_2_ signaling axis.

**Figure 6:**
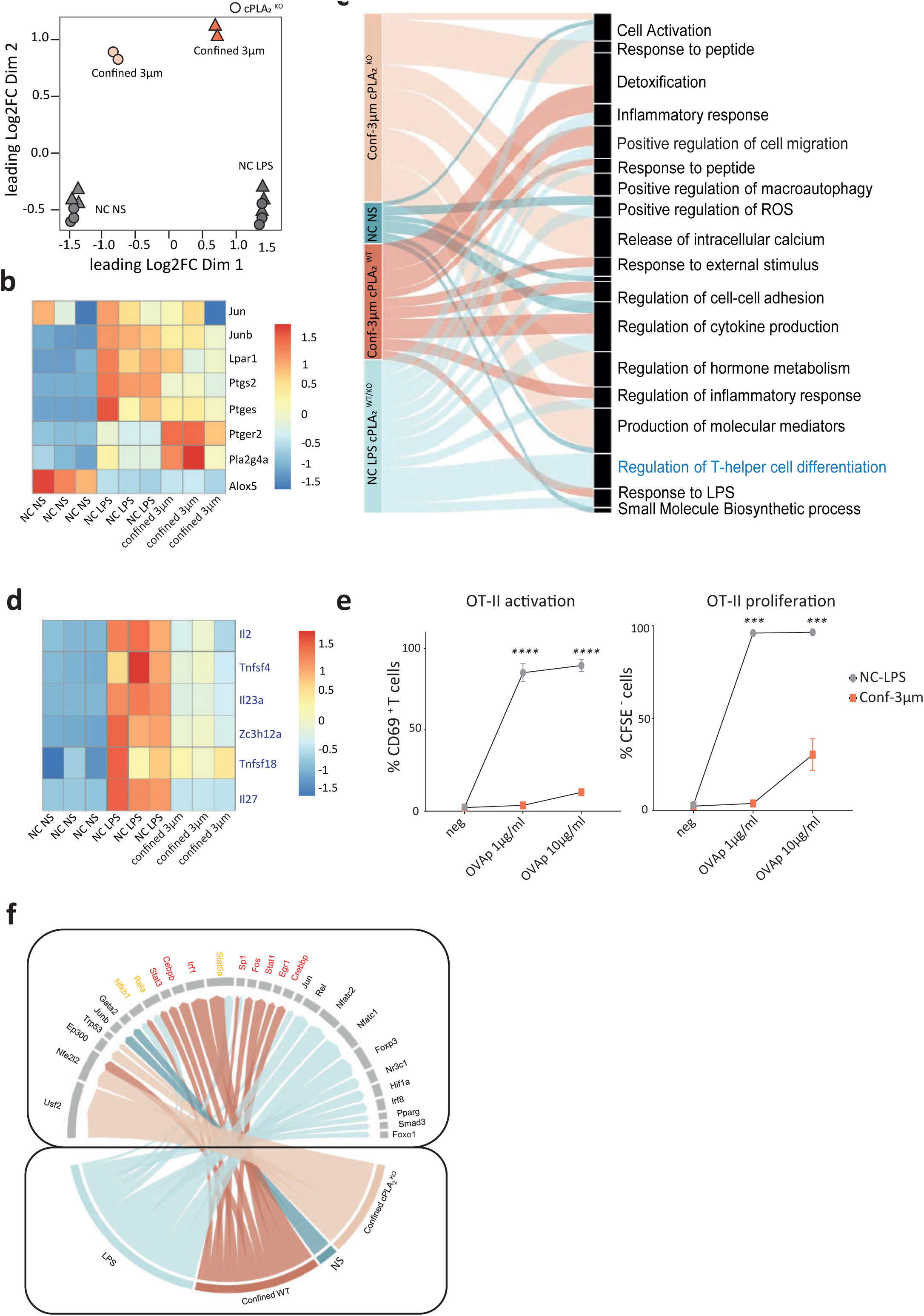
cPLA_2-_dependent transcriptional reprogramming in response to shape sensing specifically controls the immune regulatory properties of DCs. **(a)** Multidimensional scaling (MDS) of the bulk RNA-Seq sa_2_ mples (**b**) H_2_eat map of cPLA_2_-related genes including the cPLA_2_ gene itself (*Pla2g4a*) (**c**) Pathway ana_2_lysis of genes _2_differentially expressed in confined versus LPS-treated DCs (for 4h in both cases); In the alluvial plot arrow width corresponds to the enrichment score from the gene ontology analysis. All reported gene ontologies were significantly enriched (p value < 0.05) (**d**) Heat map of genes in the “regulation of T helper differentiation” pathway marked in red in figure **c** (**e**) Antigen presentation assays of OT-II (CD4) T-cells incubated with DCs either activated with LPS or confined at 3 µm-height. Left panel: percentage of CD69^+^ CD4^+^ OT-II T cells in live cells after 18 h of incubation with OVA peptide II and pre-incubated with 3 µm-confined or LPS-treated DCs for 4h. Right panel: Percentage of CFSE− CD4^+^ OT-II T cells in live cells after 3 days of incubation with OVA peptide II and pre-incubation with 3 µm-confined or LPS-treated DCs for 4h, analyzed by flow cytometry. Graph: Mean ±SEM, N=5, P values multiple t-test, ***: p = 0.0001, ****: p < 0.0001. (**f**) Transcription factor (TF) analysis on the different conditions. Transcription factor activity estimation used the TRUUST database which predicts TF activity and assigns a score. Arrow thickness corresponds to the enrichment score of each TF.

To better understand the specificity of DC reprogramming in response to shape sensing, we compared the pathways induced by confinement or by LPS treatment of DCs (fig. 6c). One of the most striking differences observed was related to the “regulation of T helper cell differentiation” pathway that was exclusively induced by LPS (fig. 6c, d). Mounting an efficient T cell response against microbial threat typically requires three signals from DCs: specific antigenic peptide presented on Major Histocompatibility Complexes class (MHC, signal 1), co-stimulation (signal 2) and cytokines (signal 3). The two latter ensure T cell activation rather than tolerization, as well as proper proliferation. Accordingly, we observed that genes encoding for MHC-II as well as key co-stimulatory molecules (CD80, CD86) were expressed at lower levels in confined DCs as compared to LPS-treated cells (Fig. S4b), leading to lower surface expression levels (of note the difference is not statistically significant) (fig. S4c). Confined DCs also express less stimulatory cytokines (IL2, IL12, IL15, IL27, fig. s4b), resulting in lower secretion levels (fig. s4d). This suggests that confined DCs might be less potent than LPS-treated cells for T lymphocyte activation. To test this hypothesis, we loaded the two types of DCs with the class-II ovalbumin (OVA) antigenic peptide and incubated them with OT-II transgenic T cells, which specifically recognize these MHC class II-peptide complexes. OT-II cell stimulation with DCs that experienced confinement led to both lower T cell activation (quantified as upregulation of CD69) and lower proliferation (quantified by CFSE dilution) than stimulation with LPS-treated cells (fig. 6e), although cell viability was comparable in both conditions (fig. S4e). These results confirmed that confined DCs were less potent than LPS-treated cells in T lymphocyte activation. Hence, DCs experiencing shape changes are in a distinct stage as compared to microbe-activated DCs, displaying a milder T cell activation capacity, consistent with the tolerogenic properties that had been previously proposed for DCs migrating to lymph nodes at steady-state (52). Altogether, our data suggest that the Arp2/3-cPLA_2_ shape sensing pathway could be unique in its capacity to tune steady-state migrating DCs to the lymph nodes and endows them with tolerogenic potential.

To gain insights into the mechanisms accounting for this specificity, we compared the transcription factor binding sites found in the promoters of the genes enriched in DCs confined at 3 µm-height or treated with LPS. Such analysis allows inferring the nature of the transcription factors differentially implicated in the two types of DC responses. We found that DC confinement leads to specific activation of IRF1-, Stat1, Stat3 and Sta5a-dependent gene transcription, which are not activated by LPS (fig. 6f). These transcription factors are known to be involved in IFN signaling and/or activation of NFκB (53), in good agreement with our results highlighting that these two pathways are enriched upon shape sensing induced by confinement (fig. 5d). Noticeably, IRF1, Stat3 and Stat5a had been implicated in acquisition of tolerogenic properties by DCs (54; 55; 56; 57; 58; 59; 60) consistent with the ARP2/3- cPLA_2_ shape sensing axis serving maintenance of peripheral tolerance. Accordingly, none of these transcription factors were activated in cPLA_2_^KO^ confined DCs (fig. 6f). Altogether our results suggest that the interplay between the cytoskeleton and the lipid metabolism enzyme cPLA_2_ transcriptionally reprograms DCs in response to precise shape changes, endowing them with the ability to reach lymph nodes in an immunoregulatory state compatible with their tolerogenic homeostatic function.

## Discussion

Here we show that DCs are equipped with a mechanism of shape sensing that defines their migratory behavior. Cell shape changes induced by confinement trigger an Arp2/3 dependent-actin polymerization in the peri-nuclear area and leads to the nuclear translocation and activation of the cPLA_2_ lipid metabolism enzyme. This in turn induces activation of NFκB and specific reprogramming of DCs, which includes the upregulation of the chemokine receptor CCR7 expression, licensing DCs for migration to lymph nodes even in the absence of external inflammatory stimuli. These findings might explain why, despite two decades of research, the signals responsible for homeostatic DC migration had remained unknown: rather than sensing (bio)chemical signals, DCs might sense the physical constrains they encounter while patrolling their environment through this Arp2/3-cPLA_2_-NFκB shape sensing pathway.

Previous work from us and others suggest that a specific threshold of nuclear deformation can lead to nuclear envelope tension and cPLA_2_ insertion into the nuclear membrane, which leads to its activation (27; 29). Both the ability of cPLA_2_ to insert into the membrane upon tension and to respond to extracellular calcium entry endow this enzyme with mechanosensing properties (61). How nuclear membranes get tensed is still largely unknown, however, drawing a parallel with the plasma membrane and the actin cortex, it is likely that nuclear membranes tension depends on forces produced by actin, and in particular branched actin (62; 63), via the LINC complex. Such forces might also open nuclear pores to promote further cPLA_2_ translocation from the cytoplasm to the nucleus, similar to what had been shown for YAP/TAZ (64). Both these scenarios are consistent with the role of Arp2/3 here described. Of note, membrane tension is more likely to rise in the inner nuclear membrane, as the outer one is continuous with the endoplasmic reticulum and thus possesses a very large membrane reservoir. Although further work will be needed to fully decipher the precise mechanism of shape sensing in DCs, we propose that it may involve Arp2/3-branched actin-dependent nuclear pore opening and tension increase of the inner nuclear membrane, leading to cPLA_2_ nuclear translocation, insertion, and activation.

We postulate that self-activation of the ARP2/3-cPLA_2_ axis originates from the successive events of deformation that DCs undergo as they move through the complex environment of peripheral tissues. These deformation events would induce a ‘cellular massage’ and thereby activate the shape sensing pathway in DCs. Notably, the activation of this shape sensing pathway is induced at a precise amplitude of cell deformation, compatible with the shape changes observed in DCs patrolling tissues such as the skin (fig 1a) (65; 25). Yet, the physical constraints imposed by the environment are likely to vary in distinct tissues (66; 67) or distinct pathological contexts, such as the tumor environment (68; 69), even if they globally stay in the same range. Determining whether the range of sensitivity of DCs to deformation is adapted to each tissue would be of the upmost importance to understand how the Arp2/3-cPLA_2_ shape sensing pathway controls both DC tissue exit to lymph nodes and their immunoregulatory properties toward T cells.

This ARP2/3-cPLA_2_ shape sensing pathway is the first one identified so far to be specifically involved in homeostatic DC migration to lymph nodes, a process essential for the maintenance of tolerance. Our results show that this shape sensing axis has no impact on reprogramming of DCs by microbial components such as LPS. Importantly, even though confined and LPS treated DCs share a considerable part of their transcriptional program, LPS treatment led to activation of specific pathways associated with efficient T cell activation, which were not induced by confinement. Consistently, confined DCs were functionally less efficient in triggering T cell activation as compared to LPS treated cells. These results are in good agreement with the tolerogenic function proposed for homeostatic DC migration to lymph nodes (49; 70). Furthermore, several of the transcription factors specifically activated in confined cells have been described as associated to immune tolerance (54; 57; 58; 71; 59). These data highlight that the cell shape changes that DCs experience while sampling antigens in peripheral tissues at steady-state endows them with immunoregulatory properties. This would enable them to transport self-antigen and reach lymph nodes in a tolerogenic state, essential for the maintenance of homeostasis. The ability of DCs to integrate both physical cues of their environment and biochemical ‘danger’ cues provided by microbial or tumoral threats and define which signal dominates would determine in which state they reach lymph nodes and the outcome of the T cell immune response they initiate. Future studies aimed at determining how this pathway is finely tuned in tissues should help understanding the contribution of the shape sensing pathway to the maintenance of tolerance and immune responses. One possibility could be that the sensitivity of the Arp2/3-cPLA_2_ shape sensing pathway in DCs is finely tuned in each tissue by the level of WASp and/or Arpin activity (73; 70).

Importantly, the here described ARP2/3-cPLA_2_ shape sensing axis could also impact the migration, function and fate of additional motile cPLA_2_-expressing cells, for example tumor cells that use CCR7 to invade healthy tissues by first migrating to lymph nodes (7; 8). Evaluating the role of this shape sensing pathway in these highly motile cells will therefore be of the highest interest to understand how they spread and invade healthy tissues.

## Acknowledgments

Funding: This project has received funding from the European Union’s Horizon 2020 research and innovation program under the Marie Skłodowska-Curie grant agreement No 666003. This work has also received support under the program «Investissements d’Avenir» launched by the French Government (ANR-10-IDEX-0001-02 PSL), from la Fondation de la Recherche Médicale (Grant SMC202006012351 to AMLD). P.J.S received the support of Human Frontier Science Program (HFSP) RGP0032-2022 and Forschungszentrum Medizintechnik Hamburg (FMTHH, grant 04fmthh2021). H.D.M received the support of ANR-20-CE15-0023 (InfEx). High-throughput sequencing was performed by the ICGex NGS platform of the Institut Curie supported by the grants ANR-10-EQPX-03 (Equipex) and ANR-10-INBS-09-08 (France Génomique Consortium) from the Agence Nationale de la Recherche (“Investissements d’Avenir” program), by the ITMO-Cancer Aviesan (Plan Cancer III) and by the SiRIC-Curie program (SiRIC Grant INCa-DGOS-465 and INCa-DGOS-Inserm_12554). Data management, quality control and primary analysis were performed by the Bioinformatics platform of the Institute Curie. We would like to also acknowledge the imaging and flow cytometry facilities in the Institute Curie for their great help and equipment.

## Material and methods

### Mice

C57BL6/J mice were obtained from Charles River, catalog #000664. CCR7-GFP knock in/ knock out mice were obtained from Jackson laboratory (stock# 027913), bred in our animal facility, original paper: Nakano H et al., 2013. CD11c-Cre mice: bred in our animal facility (Caton et al., 2007). Arpin and cPLA_2_ conditional knockout mice were generated by CIPHE (Centre d’Immunophénomique) Marseille, France. Both mice were generated using CRISPER cas-9 technique to generate cPLA_2_ and Arpin FLOX mice that were later crossed in our animal facility with Cd11c Cre mice. Ikkβ knockout mice were described in Baratin et al., 2015. WASp KO mice on a C57BL/6 (CD45.2) genetic background were from Federica Benvenuti lab mice. Experiments were performed using homozygous WASp –/– females or males as KO. Littermates or age-mated mice were used as controls for all experiments involving knockout animals; breeder mice were previously backcrossed to C57BL6 for 7 generations. CD11c-Cre+ mT/mG+ mice used for intravital imaging were previously described (Muzumdar et al. (2007).

### Cells

Dendritic cells (DCs) were obtained following a protocol first described by (K. Inaba 1992). Both whole legs from 6 to 8 weeks old mice were flushed to obtain bone marrow. Cells are maintained in culture during 10 days in IMDM medium (Sigma-Aldrich, Darmstadt, Germany) containing 10% FBS decomplemented and filtered (Biowest, Nuaillé, France), 20 mM L-glutamine (Gibco, Waltham, Massachusetts, USA), 100 U/ml penicillin–streptomycin (Gibco, Waltham, Massachusetts, USA), 50 μM 2-mercaptoethanol (Gibco, Waltham, Massachusetts, USA), and 50 ng/ml of GM-CSF containing supernatant obtained from transfected J558 cells tested by ELISA, as previously described by (Faure-Andre 2008). At days 4 and 7 of culture, cells are detached using PBS-EDTA (5 mM) and replated at 0.5:10^6^ cells per milliliter of medium. At day 10, 90% of adherent cells express CD11c, an integrin family member, as well as MHC class II at medium levels which is specific to DCs (36). The obtained cells can be used at day 10 or 11 as immature cells. The generation of DCs from bone marrow supplemented with GM-CSF is a well-defined protocol but it promotes the differentiation of three cell types: granulocytes, macrophages, and DCs (74) but they can be separated during the cell culture based on their adhesion. Granulocytes are eliminated during the culture since they are non-adherent, whereas macrophages are much more adherent than the other two cell types and stick to the bottom of the plate and thus we recover the semi-adherent DCs at the last day of culture.

#### DC maturation

when using mature DCs (mainly as a positive control for maturation profile of DCs): Day 10 cells (which are immature when we recover them), are stimulated with 100 ng/ml of lipopolysaccharide (LPS) (Salmonella enterica serotype typhimurium; Sigma, Darmstadt, Germany) for 25 min and washed three times with complete medium, re-plated in fresh medium, left overnight in the incubator and then used. LPS-activated DCs have higher levels of expression of the costimulatory molecules CD86 and CD40 as well as the chemokine receptor CCR7.

### 6-well plate confiner

Dendritic cell confinement was performed using a 6-well plate confiner described by (Y.-J. Liu 2015) that allows the recovery of large number of cells and the simultaneous imaging of several conditions. To make the polydimethylsiloxane (PDMS, RTV615) pillars at a certain height, 12 mm glass coverslips were sonicated in methanol, washed in ethanol then plasma treated and placed on the top of a PDMS mixture on top of wafer molds that contain the pillars at the desired height. The PDMS mixture is composed of PDMS A/crosslinker B at 1/10 W/W. The height of the PDMS pillars is what determines the height of confinement of the cells between the coverslip and the bottom substrate. After adding the coverslips to the wafers, they are baked at 95° for 15 min, then carefully removed using isopropanol. Then they are washed again with isopropanol, dried well and plasma treated for 2 min. They are then incubated with non-adhesive pLL-PEG (SuSoS, PLL (20)-g [3.5]-PEG (2)) at 0.5mg/ml in 10mM HEPES pH 7.4 buffer for one hour at room temperature. The coverslips are then washed well with water to remove all the remaining PEG and incubated in cell’s medium for at least 2 h before the confinement starts. To make the confinement steps, we use a modified version of a classical 6-well plate cover; large PDMS pillars were stuck on the coverlid (these are the ones which will hold the fabricated coverslips). These large pillars will push the coverslips from the top of the lid to confine the cells. 6-well plates with glass bottom can be used in case of imaging of the cells (MatTek corporation, P06G-1.5-20-F).

### Live cell imaging

Live Time-lapse recordings were acquired with 20x (NA 0.75) dry objective, Nikon video-microscopy, for 4-6 h at 37 °C with 5% CO2 atmosphere. Or by confocal microscope (Leica DMi8, SP8 scanning head unit) with 40x (NA 1.3) oil objective with a resolution of 1024×1024 pixels. Both microscopes were controlled by Meta Morphe software. Image analysis was performed using Imag eJ software (75) (NIH, http://rsb.info.nih.gov/ij/index.html). For GFP quantification, imaging was done using both transmission phase and GFP. Images corresponding to each time point of interest were collected in each condition, the outline of each cell was drawn by hand on the trans images, then the mean GFP signal was calculated on the corresponding GFP image using a homemade macro; mean intensity of each cell was subtracted from the average intensity of the background, then multiplied by cell’s area to calculate total GFP intensity per cell. Of note, only cells with intensity higher than the background were plotted (percentage of cells is indicated in the legends).

### Constructs

NLS_GFP: pTRIP-SFFV-EGFP-NLS (NLS-GFP hereafter) was generated introducing the SV40 NLS sequence (PKKKRKVEDP) by overlapping PCR at the C-terminal of GFP in pTRIP-SFFV.

### Lentivirus transduction

Transduced DCs were obtained by transfection of BMDCs from C57BL/6 mice were 1million cells/2ml medium were plated. At day 4, 40ml of fresh pTRIP-SFFV-GFP-NLS lentivector supernatant were loaded in Ultra-Clear Centrifuge tubes (Beckman Coulter) and ultracentrifuged at 100,000g in a SW32 rotor (Beckman coulter) for 90 min at 4°C and re-suspended in 400µl of in DC medium. 200µl of ultracentrifuged virus were used to infect one well of cells in presence of 8µg/ml of Protamine. Cells were then left for 48h, then washed to remove the viral particles and left in new medium till day 10 of culture.

### Drugs and Reagents

For live imaging experiments: NucBlue (Hoechst33342) from Thermo fischer (R37605) to mark the DNA. For drug treatments: AACOCF3 (selective phospholipase A_2_ inhibitor, Tocris #1462-5), CK666 (selective Arp2/3 inhibitor, Tocris #3950), BI605906 (selective IKKβ inhibitor #53001).

### Immunofluorescence microscopy

The staining was done directly on the confinement plate to not affect the expression of the markers of interest. After removing the confinement led, samples were fixed directly with 4% paraformaldehyde for half an hour, permeabilized with 0.2% Triton X-100, and then incubated overnight with the primary antibodies at 4°. The next day, samples were washed with PBS and incubated with the corresponding secondary antibodies for one hour, then washed three times with PBS and mounted with fluromount solution. Imaging was done using confocal microscope (Leica DMi8, SP8 scanning head unit) with 40x (NA 1.3) oil objective with a resolution of 1024×1024 pixels. The following primary antibodies were used for the IF staining: anti-CCR7 (abcam #ab32527), anti-laminA/C (Sigma #SAB4200236), anti-cPLA_2_ (abcam #ab58375), anti-NF-κB P65 (cell signalling #mAB 8242), Alexa Fluor-coupled Phalloidin (Invitrogen).

#### Calculation of the fluorescence intensity

Images were acquired with SP8 confocal microscope, then Z stacks of usually 0.33 µm as a step size were made for each position, with a home-made macro, the plan of the nucleus was used to quantify the florescence intensity by making a mask on both the nucleus (taking the Dapi channel) and the cell (taking the phalloidin channel) and intensity was calculated.

### Transfection and SiRNA

Bone marrow derived DCs at day 7 (3×10^6^) were transfected with 100μl of the Amaxa solution (Lonza) containing siRNA (control or target-specific) following the manufacturer’s protocol. Cells were further cultured for 48-72 h. At the day of the experiment wells were detached with medium, some cells were collected to check the efficacy of the Si step by either western blot or RT-qPCR protocols. The following SMARTpool siRNAs were used: SMARTpool: ON-TARGETplus *Pla_2_g4a* siRNA (Dharmacon # L-009886- 00-0010) and ON-TARGETplus Non-Targeting Control Pool (Dharmacon # D-001810-10-20).

### RT-qPCR

After removing the confinement, the lysis buffer was added directly to recover the confined cells, taking in parallel control cells that were not confined. RNA extraction was performed using RNeasy Micro RNA kit (Qiagen), according to the manufacturer’s protocol. cDNA was produced using the high-capacity cDNA synthesis kit (thermo fisher), according to the manufacturer’s protocol, starting from 1 μg of RNA. Quantitative PCR experiments were performed using Taqman Gene Expression Assay (Applied Biosystems) and carried out on a Lightcycler 480 (Roche) using the settings recommended by the manufacturer. The following primers were used: Mm99999130_s1 for CCR7, Mm01284324_m1 for Pla_2_g4a and Mm99999915 for GAPDH as a control. The expression of each gene of interest was assessed in immature non-confined cells. Samples were run in triplicate for each condition. Data were subsequently normalized to GAPDH values, and to the values obtained in control immature cells were used as a base unit equal to one, thus allowing for display of the data as “fold-greater” than the immature cells. The fold change was calculated by the formula 2^- ΔΔCT^.

### Flow cytometry

To characterize the expression of some surface markers on the DCs after confinement we used the FACS approach. In brief, after confining the cells, the confiner was removed, and cells were recovered directly by gently washing with medium. Non-confined cells activated with LPS were also taken as controls. Cells were re-suspended in the buffer (PBS BSA 1% EDTA 2mM). After blocking with Fc antibody (BD #553142) and live/dead staining kit (Thermo # L34966) for 15 min, cells were stained with the desired antibodies for 20 min (at 37° for CCR7 staining and at 4° for the rest). Cells were then washed three times and re-suspended in the staining buffer. Flow cytometry was performed on LSRII (BD) and analyzed using FlowJo software version 10. Mean florescent values of each condition were plotted with Graphpad Prism version 8. The following antibodies were used: anti-CD80 (BD # 553769), anti-MHCII (Ozym # BLE107622), anti-CD86 (Ozyme # BLE105037), anti-CD11c (BD # 550261) with the corresponding isotypes to each antibody.

#### For the T-cell presentation assay

OVA peptide (Cayla #vac-isq) was added to each condition with the corresponding concentration, cells were then confined for 4 h, then recovered by washing with medium and counted. T-cells which were purified from OT-II mice, stained with CFSE and added to the recovered DCs with a ratio of 10 to 1 respectively. Cells were plated in round bottom 96 well plates. OT-II T cell activation was analyzed 18 h after, and after 3 days, proliferation was measured by flow cytometry. Cells were stained in 2mM EDTA, 5%FBS in PBS following the same protocol as before. Then 2 days later, the rest of the cells were recovered and analysed for their proliferation using CFSE. Flow cytometry was performed on LSRII (BD) and analyzed using FlowJo software version 10. Percentage values were plotted with Graphpad Prism version 8. The following antibodies were used: anti-CD4 (BD # 553051), anti-CD69 (eBioscience # 48-0691-82), anti-TCR (BD # 553190).

### RNA seq

After removing the confinement, the lysis buffer was added directly to recover the confined cells, taking in parallel control cells that were not confined. RNA extraction was performed using RNeasy Micro RNA kit (Qiagen), according to the manufacturer’s protocol. The samples were checked for the quality of the extracted RNA before sending them to the NGS-sequencing platform in the institute Curie RNA sequencing libraries were prepared from 300ng to 1µg of total RNA using the Illumina TruSeq Stranded mRNA Library preparation kit and the Illumina Stranded mRNA Prep Ligation kit which allow to perform a strand specific RNA sequencing. A first step of polyA selection using magnetic beads is done to focus sequencing on polyadenylated transcripts. After fragmentation, cDNA synthesis was performed and resulting fragments were used for dA-tailing and then ligated to the TruSeq indexed adapters (for the TruSeq kit) or RNA Index Anchors (for the mRNA Ligation kit). PCR amplification was finally achieved to create the final indexed cDNA libraries (with 13 cycles). Individual library quantification and quality assessment was performed using Qubit fluorometric assay (Invitrogen) with dsDNA HS (High Sensitivity) Assay Kit and LabChip GX Touch using a High Sensitivity DNA chip (Perkin Elmer). Libraries were then equimolarly pooled and quantified by qPCR using the KAPA library quantification kit (Roche). Sequencing was carried out on the NovaSeq 6000 instrument from Illumina using paired-end 2 x 100 bp, to obtain around 30 million clusters (60 million raw paired-end reads) per sample.

### RNA seq data analysis

Raw data are available at GEO under the number: GSE207653.

#### Analysis of sequencing data quality

reads repartition (e.g., for potential ribosomal contamination), inner distance size estimation, gene body coverage, strand-specificity of library were performed using FastQC, Picard-Tools, Samtools, and RSeQC. Reads were mapped using STAR [PMID: 23104886] on the mm39 genome assembly.

Gene expression was estimated as described previously [PMID: 34495298] using Mouse FAST DB v2021_2 annotations. Only genes expressed in at least one of the two compared conditions were analyzed further. Genes were considered as expressed if their Fragments Per Kilobase Million (fpkm) value was greater than fpkm of 98% of the intergenic regions (background). Analysis at the gene level was performed using Deseq2[PMID: 25516281]. Genes were considered differentially expressed for fold-changes ≥1.5 and p-values ≤0.05. Pathway analyses and transcription factor network analysis were performed using WebGestalt 0.4.4 (76)[PMID: 31114916] merging results from upregulated and downregulated genes only, as well as all regulated genes. Pathways and networks were considered significant with a false discovery rate ≤0.05. The graphics (heat map, MDS, scatter and volcano plots) were generated using R v4.2.1 with the help of pheatmap, ggplot2, and EnhancedVolcano [PMID:27207943] packages respectively. The heat maps were created using the Z-score of EdgeR normalized counts.

### Method for analysis of microarray GSE49358

The GSE49358 microarray data (Tamoutounour et al., 2013) were downloaded from GEO. Microarray data were annotated using the ‘mogene10stprobeset.db’ R package (v. 2.7). cPLA signature genes were displayed as heatmaps using tidyverse (v. 1.3.0), scales (v. 1.1.1), readxl (v. 1.3.1).

### In vivo analysis of DC subsets

Littermates at the age of 8-12 weeks of different WT and KO mice were used. Both males and females. To collect cells from the skin, 1 cm^2^ part of the skin was cut with the help of a stencil and transferred to an Eppendorf tube. 1 ml of 0.25 mg/ml of liberase (Sigma #5401020001) and 0.5 mg/ml of DNAse (Sigma #10104159001) in 1 ml RPMI medium (sigma). The skin was cut using scissors and incubated 2 h at 37°. To collect cells from lymph nodes, inguinal lymph nodes were removed. Then they were transferred to an Eppendorf tube containing 500 µl of RPMI medium, DNAse was added at concentration of 0.5 mg/ml and collagenase at 1 mg/ml. The lymph nodes were further cut with scissors and incubated 20 min at 37°. Then, cells were re-suspended in PBS 0.5% BSA 2 mM EDTA at 4°C. Cells were filtered and then stained following the same protocol as explained before. The following antibodies were used: Anti-CCR7 (Biolegend #120114), anti-CD11c (eBioscience #25-0114- 81), anti-CD326 (Biolegend #118217), anti-CD86 (BD pharmingen #553692), anti-CD11b (Biolegend #101237), anti-MHCII (eBioscience #56-5321-80), anti-CD45 (eBioscience #61-0451-82), anti-CD8a (BD bioscience #553035), anti-CD103 (eBioscience #46-1031-80). Counting beads were used to normalize the number of cells to the number of beads. Flow cytometry was performed on LSRII (BD) and analyzed using FlowJo software version 10. Percentage values were plotted with Graphpad Prism version 8.

### Intravital imaging of DC migration in ear skin dermis by 2-photon live-imaging

Data previously generated (Moreau et al, Dev Cell 2019) were reanalyzed to estimate the deformation experienced by DCs migrating in the skin. Two-photon intravital imaging of the ear dermis was performed as previously described (Filipe-Santos et al., 2009), on bone-marrow chimeras reconstituted with 50% of CD74WT bone marrow (from mTmGflox/flox CD11c-Cre+ animals: DCs are GFP+) and 50% CD74^KO^ bone marrow (from CD11c-EYFP animals: DCs are YFP+). Imaging was done at least 8 weeks after irradiation (9Gy) and bone-marrow reconstitution to ensure steady-state. In brief, mice were anesthetized and placed on a custom-designed heated stage and a coverslip sealed to a surrounding parafilm blanket was placed on the ear, to immerge a heated 25X/1.05 NA dipping objective (Olympus). Imaging was performed using an upright FVMPE-RS microscope (Olympus). Multiphoton excitation was provided by an Insight DS + Dual laser (Spectra-Physics) tuned at 950 nm. Emitted fluorescence was split with 520, 562 and 506 nm dichroic mirrors and passed through 593/40 (mTom) and 542/27 (YFP) filters to nondescanned detectors (Olympus) and 483/32 (collagen by second harmonic generation) and 520/35 (GFP) filters to GASP detectors (Olympus). Typically, images from about 10 z planes, spaced 4 μm were collected every 45 seconds for up to one hour. We focused our new analysis of these existing movies on the GFP^+^ cells which are CD74^WT^ skin DCs.

### Statistics

Statistical significance was calculated between two groups by Mann-Whitney test. Ordinary Mann-Whitney with multiple comparison was used to calculate statistical significance between multiple groups. Analyses were performed using GraphPad Prism 8 software.

## Supplementary figures

**Figure S1:**
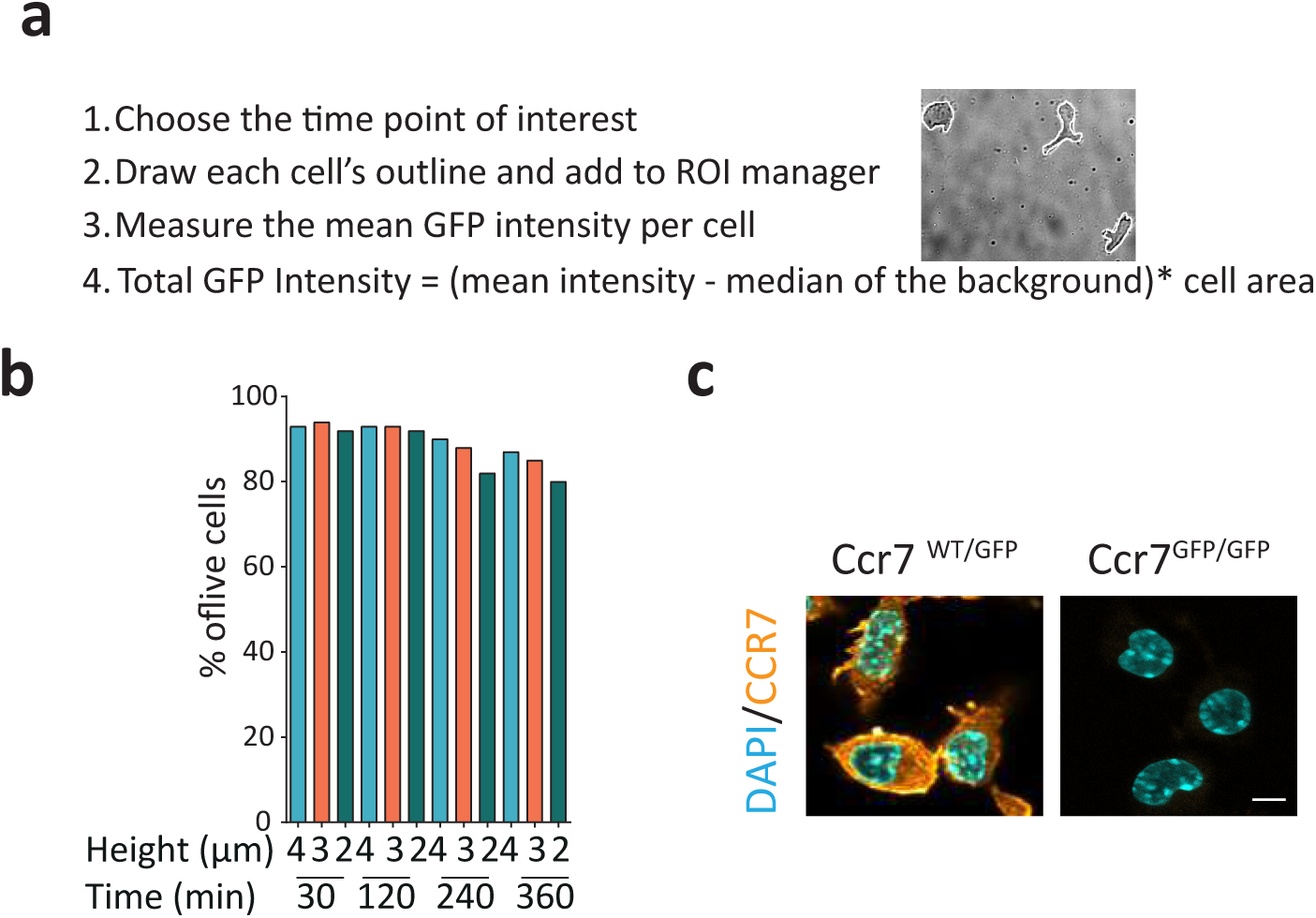
GFP quantification approach. (**a**) Quantification approach to quantify GFP intensity in each cells. (**b**) Quantification of cell viability in different confinement conditions and during different time points, using propidium iodide (**c**) representative images of immature DCs confined for 4h at the height of 3 µm from C57BL/6J (WT) or GFP/GFP mice, CCR7was visualized using immunostaining (in orange) and the nucleus stained with Dapi (in cyan) scale bar10 µm.

**Figure S2:**
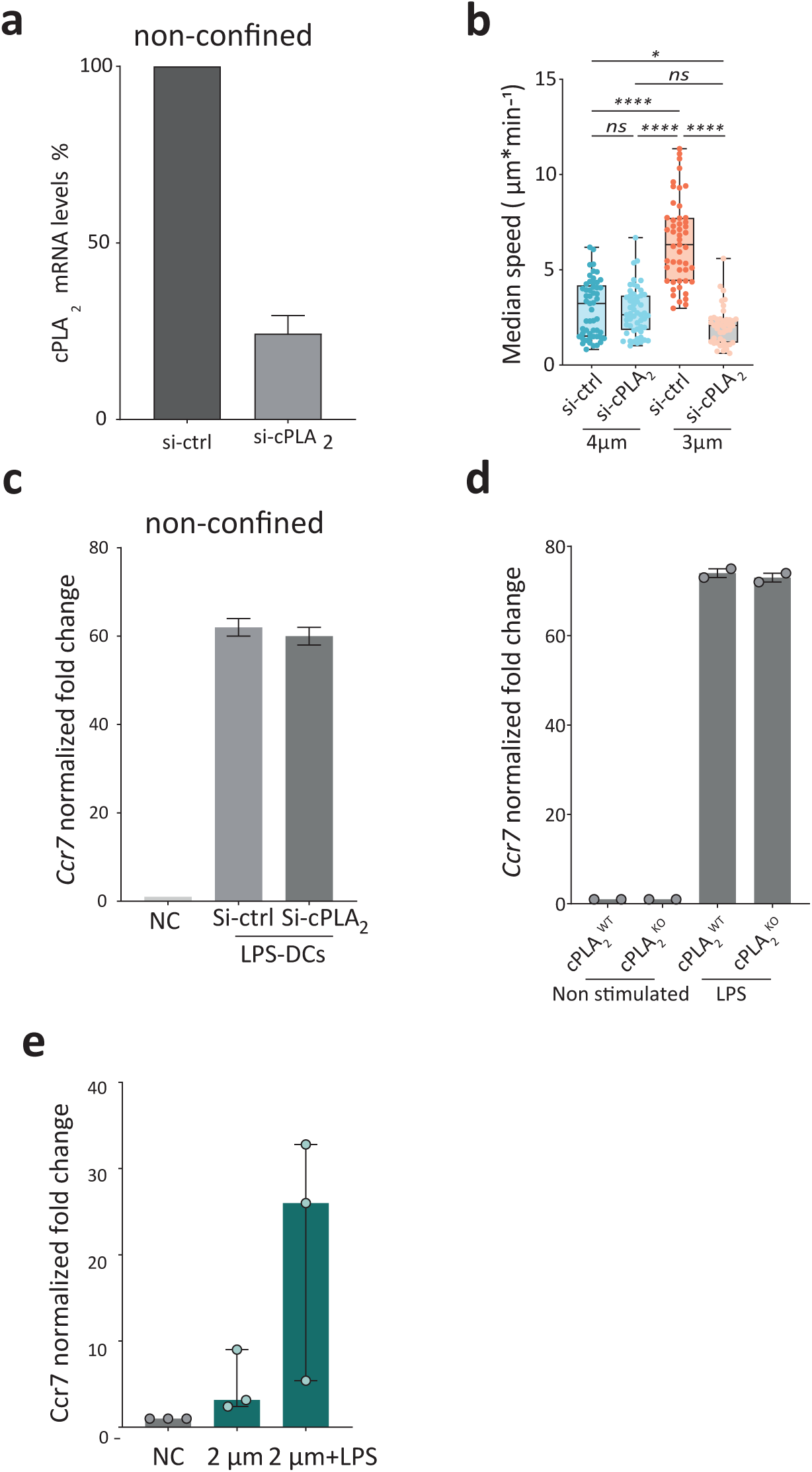
cPLA2 activity is not necessary for LPS-induced *Ccr7* up regulation upon confinement. (a) RT-qPCR data of cPLA_2_ gene expression upon knock down of cPLA_2_ with Si-RNA technique, N=3. (b) Median speed of cPLA_2_ KD or control DCs confined at 4, and 3µm. N=3, n=53 cells in cPLA_2_ Ctrl confined at 4µm, n= 53 cells in cPLA_2_ KD confined at 4µm, n=49 cells in cPLA_2_ Ctrl confined at 3µm, n=44 cells in cPLA_2_ KD confined at 3µm. P value Kruskal-Wallis test, ****: <0.0001, *: 0.0315, ns: non-significant. (**c**) RT-qPCR data of *Ccr7* gene expression in cPLA_2_ KD or control DCs activated with LPS, data shows no difference in *Ccr7* up regulation in response to LPS in both cells types. N=2. (**d**) RT-qPCR data of *Ccr7* gene expression in cPLA_2_ WT and KO DCs activated with LPS, data shows no difference in *Ccr7* up regulation in response to LPS in both cells types. N=2. (**e**) RT-qPCR data of *Ccr7* gene expression in cells confined at 2 µm height and cells activated with LPS after confinement at 2 µm height, N=2.

**Figure S3:**
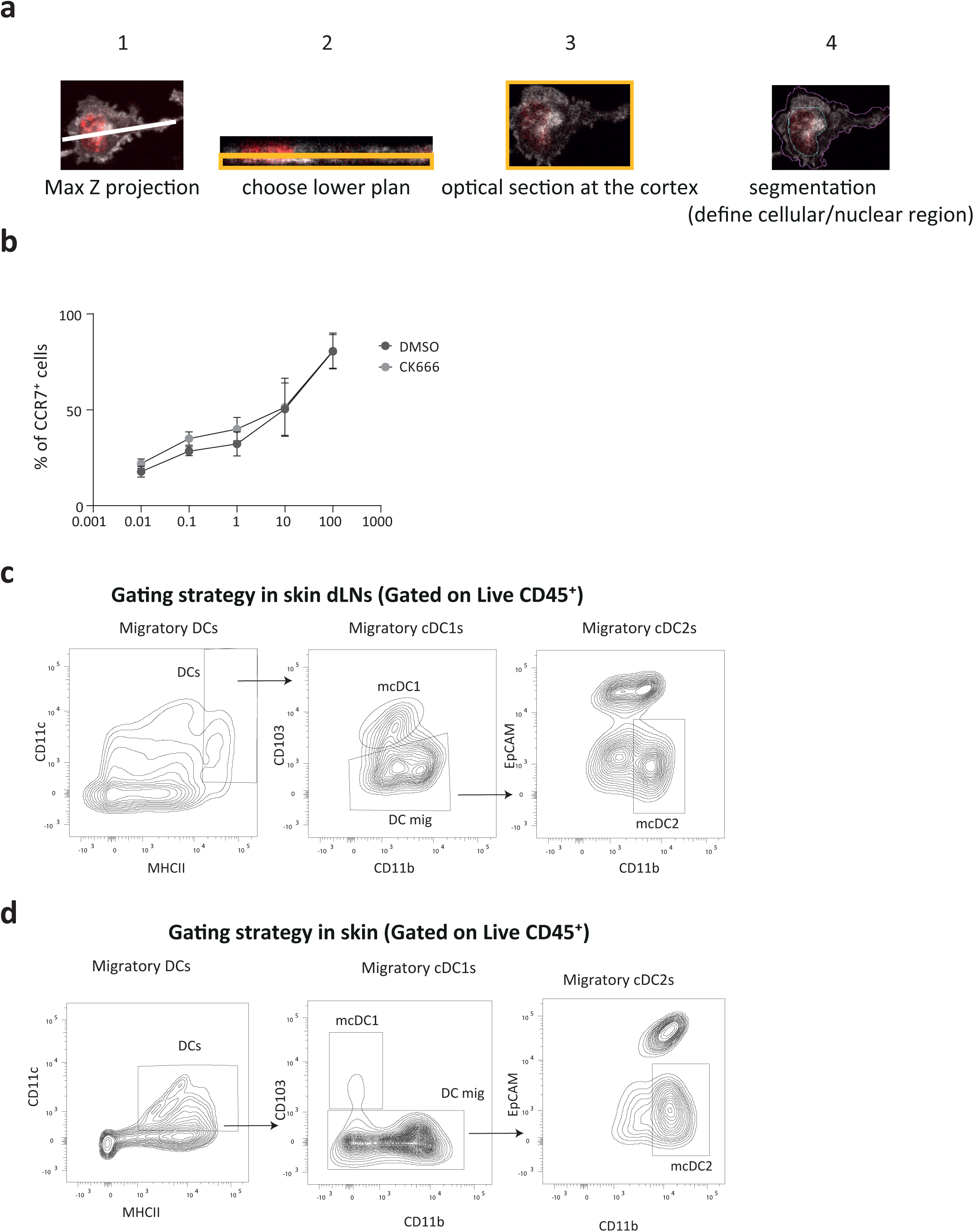
Arp2/3 branched actin is not important for CCR7 upregulation in LPS. (**a**) Quantification approach of LifeAct-GFP intensity in live cells underconfinement: 1- choose Z-stack lower plan (since cells don’t always touch the upper plan). 2- optical section at the surface cortex. 3-Segmentation to define cell and nuclear contour at the surface. 4- Measurement of LifeAct-GFP ratio: Actin Ratio= Nuclear surface mean actin intensity/cell surface meanactin intensity (ratio <1 => actin is mostly cortical). (**b**) FACS analysis of CCR7^+^ DCs activated by LPS and treated with CK666 (25µM) or DMSO. N=3.(**c,d**) Gating strategy to quantify DCs in skin-draining lymph nodes and skin of mice at steady-state: after gating on live cells, immunecells were identified as CD45^high^; CD11c and MHCII were then used to differentiate lymph node-resident DCs (MHCII^low^, CD11c^high^) from migratory DCs (MHCII^high^, CD11c^high^). Among migratory DCs, mDC1s were identified as CD11b^high^, CD103^low^ and mDC2s as CD11b^high^, EPCAM^low^.

**Figure S4:**
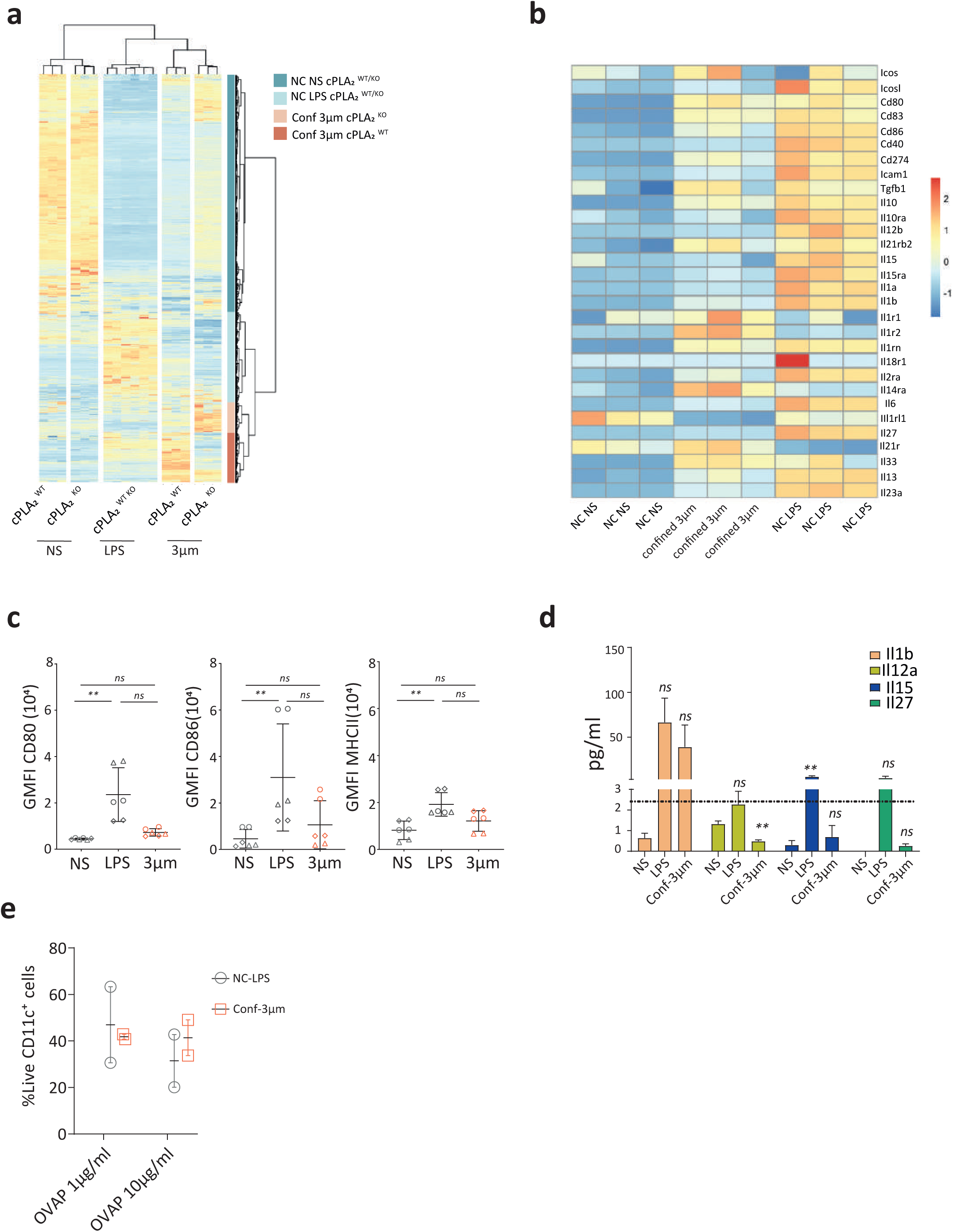
Bulk RNA-seq analysis shows transcriptional changes in cells in response to confinement in a cPLA_2_-dependent manner. (a) Heatmap of all the differentially expressed genes in cPLA_2_ WT and KO cells in all the different conditions (**b**) Heat map of examples of cytokines and costimulatory gens in DCs in response to confinement at 3 µm-height or LPS stimulation (**c**) FACS analysis of some immune-activating genes expressed_2_ by DCs not-_2_stimulated (NS) controls or is response to LPS or confinement at 3 µm h_2_eight. Gra_2_phs showing geometric mean intensityof CD80, CD86, and MHCII. N=3 each condition was done in duplicates, P value Kruskal-Wallis test (**d**) cytokine secretion analysis by luminex of some cytokines showing their protein level of expression in the supernatant of LPS or confined DCs 48h after the confinement or the LPS activation. N=5 (**e**) FACS analysis to measure the percentage of live DCs CD11c high after 48 hour of incubation with OTII T-cells shows no difference between cells either activated with LPS or confined at 3 µm in the same presentation assay experiments showed in figure 6. Graph: mean with SEM, N=2

## Supplementary movies

**Movie S1: 2 photon movie of cd11c YFP DCs in mouse ear skin.**

Example of a cd11c-YFP DCs in green migrating in ear dermis with collagen fibers observed in grey. The contour of the cells is shown in pink and arrows appear to indicate thinning events experienced by the cell while migrating. These thinning events could range in size from 2 to 5 µm. Time interval of 45 seconds and scale bar is 20 µm.

**Movie S2: Movie of BMDCs from CCR7/GFP mice under confinement.**

Bone marrow derived DCs from CCR7/GFP mice under confinement of 4, 3, and 2 µm height. Upper part is the images of cells in transmission light (in grey) and GFP signal (in green). Lower part isthe GFP signal using false colors (Physics LUT). The warmer the color the more the intensity of GFP signal. The transmission images were corrected for bleach (with the function bleach correction in ImageJ), GFP signal was smoothed using the Median filter. Time is showed in minutes (using time stamper function). Movie created with ImageJ software [67]

**Movie S3: Movie of BMDCs transduced with NLS-GFP under confinement of 2 µm height.**

Example of NLS-GFP expression in BMDCs under 2 µm height. The movie shows examples from 2 different positions of the same condition displaying rupture (when the GFP signal is diffused) and repair(when the signal is concentrated in the nucleus) events. The stacks were merged using the stack-mergefunction with ImageJ software. The movie is a Max Z projection of the GFP signal. Time is showed in minutes (using time stamper function).

**Movie S4: Movie of live BMDCs expressing LifeAct/GFP under confinement of 4 and 3 µm height.** Example of immature DCs expressing LifeAct/GFP under confinement of 4 or 3 µm, cells were stained with NucBlue to visualize the nucleus (DNA). Movies in Z-stack of 10 steps (step size of 0.33 µm).

